# Astrocyte-secreted chordin-like 1 regulates spine density after ischemic stroke

**DOI:** 10.1101/2020.12.19.423621

**Authors:** Elena Blanco-Suarez, Nicola J Allen

**Affiliations:** Molecular Neurobiology Laboratory, Salk Institute for Biological Studies, 10010 N Torrey Pines Road, La Jolla, CA, 92037, USA; Department of Neurosurgery, Thomas Jefferson University, 900 Walnut Street, Philadelphia, PA 19107, USA

**Keywords:** Astrocyte, photothrombosis, spine density, stroke

## Abstract

Ischemic stroke occurs when the brain is deprived of blood flow, preventing cells from receiving nutrients necessary to perform basic vital functions. In the peri-infarct area neurons undergo an acute loss of dendritic spines along with morphological alterations, which ultimately modify synaptic plasticity and determine neuronal survival. Astrocytes have been shown to play protective or detrimental roles in neuronal survival post-stroke, depending on the specific stage, yet we lack a clear understanding of the underlying mechanisms triggered at these different time points. Recently chordin-like 1 (Chrdl1) was identified as an astrocyte-secreted protein that promotes synaptic maturation and limits experience-dependent plasticity in the mouse visual cortex, leading us to ask if Chrdl1 regulates spine density and recovery from stroke. Using photothrombosis to model ischemic stroke, we studied Chrdl1 KO mice during the acute and subacute phases post-stroke (1 and 7 days after injury, respectively) to assess the potential of Chrdl1 to regulate spine density, glial reactivity and injury volume, characteristics that are involved in functional recovery after ischemia. We find that the absence of Chrdl1 prevents ischemia-induced spine loss in the peri-infarct area, a feature that indicates an important role of astrocytes in recovery from ischemic stroke.

## Introduction

Focal ischemic stroke is a condition characterized by the interruption of blood flow to a specific region of the brain. This interruption causes an acute depletion of glucose and oxygen, preventing cellular metabolic function, leading to cell death and tissue damage, and eventual loss of brain function (1). The brain region depleted of blood flow is referred to as the core of the injury. Mostly, the cells confined to the core of the injury follow apoptotic and necrotic pathways causing cell death and preventing neuronal rescue. The surrounding tissue suffers from a decreased blood supply, and it is called the peri-infarct area. In this region, diverse cellular and molecular mechanisms are triggered in response to the reduced blood supply and cell metabolism, which will dictate whether cells in the peri-infarct area survive or follow delayed cell death (2). It is the peri-infarct area that holds potential for recovery, and understanding the molecular mechanisms that take place in this region post-stroke is crucial to promote cell survival over delayed cell death.

Several methods are available to mimic ischemic stroke in mouse models (3). In the present study we use photothrombosis, a technique that consists of transcranial illumination of a brain region upon injection with Rose Bengal, a photoactivatable dye. This induces the formation of singlet oxygen that damages the endothelium, promoting platelet aggregation and formation of thrombi that block the blood supply to the illuminated area, causing similar effects to focal ischemia in humans (4). After ischemic stroke, there are different phases characterized by the ability for spontaneous recovery in the peri-infarct area. In the mouse, these phases are divided into acute (0-2 days post-stroke), sub-acute (2-30 days post-stroke) and chronic (beyond 30 days post-stroke) (5), with comparable time windows for humans (6). During the acute phase there is a high rate of cell death in combination with the onset of inflammatory mechanisms and formation of a glial scar. This is followed by the sub-acute phase, the time window when endogenous plasticity mechanisms take place to support neural repair. During the sub-acute phase there is a certain degree of tissue reorganization that determines the level of repair displayed during the chronic phase. At this late chronic stage, endogenous mechanisms of plasticity are diminished and neural repair is minimal (5). Currently, it is unclear what triggers each phase post-stroke, and how to harness the potential of endogenous plasticity mechanisms in order to promote functional recovery and neural repair at later stages (i.e chronic phase).

Astrocytes play important roles in homeostasis of the central nervous system (CNS), and some of these functions are of particular importance in the context of ischemic stroke, pointing to astrocytes as potential targets for neuroprotection (7). Astrocytes are important components of the neurovascular unit contributing to the integrity of the blood-brain barrier (BBB); they play an important role in glial scar formation; and they are responsible for clearing excessive glutamate from the extracellular space which otherwise triggers excitotoxic events (8). In addition, some evidence has shown how astrocytes and microglia may be involved in diverse CNS disorders and pathologies, including ischemic stroke (9, 10). Astrocytes, in response to injury and disease, go through changes in their morphology, molecular mechanisms and functionality, a response termed astrogliosis, regulated by signaling pathways that are yet to be fully defined (11). Astrocytes are not isolated, and in fact interact with a variety of cells that surround the peri-infarct area to regulate different responses such as inflammation or tissue replacement (12). Microglia are crucial in the regulation of the immune response in the CNS during stroke. They undergo morphological changes and they are recruited to the injury site where they release a series of proinflammatory molecules (13). Both astrocytes and microglia partake in inflammatory mechanisms during ischemic stroke, which promotes both detrimental and beneficial effects (14).

In the healthy brain astrocytes modulate the development, maturation and function of synapses through the secretion of diverse factors (15), and some of these factors have been linked to neuroprotective mechanisms in ischemic stroke. For example, thrombospondins-1 and -2 (TSP-1 and -2) are astrocyte-secreted proteins involved in the formation of silent synapses(16). Elimination of TSP-1 and -2 leads to a greater loss of synapses in response to ischemic stroke which impairs functional recovery, therefore establishing an important role for astrocyte-secreted proteins in synaptic regulation after stroke (17). In our previous study (18) we identified chordin-like 1 (Chrdl1), which is enriched in expression in astrocytes in upper layers of the cortex and the striatum, as an astrocyte-secreted protein responsible for synaptic maturation in the mouse visual cortex, and therefore, an important regulator of synaptic plasticity. During normal development the critical period is a time when endogenous plasticity is enhanced, permitting remodeling of visual circuits in response to a modification in visual input (19). This endogenous plasticity is greatly diminished at older ages. We demonstrated that the absence of Chrdl1 increases experience-dependent plasticity in a visual sensory deprivation paradigm, an effect that was observed not only during periods of high endogenous plasticity i.e. the visual critical period, but beyond this time into adulthood (18). A study blocking a different plasticity-limiting molecule that is expressed by neurons, PirB, found this manipulation also promoted enhanced experience-dependent plasticity, and also proved beneficial in the context of ischemic stroke by reducing the size of the injury, improving motor recovery, and decreasing astrocyte reactivity (20). This suggests that targeting endogenous plasticity-limiting molecules may be beneficial in recovery from ischemic stroke.

To enable functional recovery after ischemic stroke, plasticity potential is necessary to promote functional and structural changes to compensate for lost synapses (21). As Chrdl1 regulates synaptic maturation and limits plasticity, we therefore asked if absence of Chrdl1 is beneficial to recovery after ischemic stroke by creating a permissive environment to allow synaptic remodeling. We found that Chrdl1 is significantly upregulated during the acute phase (24 hours post-stroke) in both the peri-infarct and contralateral hemispheres, whereas at later timepoints during the sub-acute phase (7 days post-stroke) the expression of Chrdl1 remains elevated only in the peri-infarct area. Elimination of Chrdl1 *in vivo* prevents the characteristic spine loss associated with ischemic stroke that has been previously reported (22). Therefore, we hypothesize that the increase in plasticity-potential in the absence of the astrocyte-secreted protein Chrdl1 permits faster remodeling in the peri-infarct area after ischemic stroke, potentially promoting functional recovery.

## Material and Methods

### Animals

All animal work was approved by the Institutional Animal Care and Use Committee (IACUC) of the Salk Institute for Biological Studies, and the ARRIVE guidelines were followed in all animal experiments.

Mice were housed in the Salk Institute animal facility at a light cycle of 12h light:12h dark, and access to water and food ad libitum.

Wild-type (WT) mice (C57BL/6J, Jax stock number 000664) were used for analysis of expression of Chrdl1 by in situ hybridization as described below, and to breed to Chrdl1 KO mice and Thy1-YFP-J mice (B6.Cg-Tg(Thy1-YFP)HJrs/J, Jax stock number 003782). For detailed procedure to generate Chrdl1 KO mice see reference (18). For the generation of experimental mice heterozygous (+/-) Chrdl1 females (as Chrdl1 is in the X chromosome) were bred to WT (+/y) C57BL/6J males. All experiments were performed using male Chrdl1 KO (-/y) and WT (+/y) littermates. Thy1-YFP-J male mice were crossed with heterozygous Chrdl1 KO (+/-) females for the generation of Thy1-YFP-J Chrdl1 KO (-/y) and WT (+/y) males used for experiments (referred to as YFP-Chrdl1 KO and WT).

### Photothrombotic ischemic stroke

Male mice at 4 months of age were placed on a stereotaxic frame and anesthetized by 2% isofluorane in oxygen by constant flow via nose cone. Mice were retro-orbitally injected with 10 mg/ml Rose Bengal (Fisher R323-25) in saline (0.9% NaCl) at a dose of 25mg/kg. Rose Bengal was prepared fresh before surgeries and protected from light. Control mice subjected to sham surgeries were injected with the equivalent volume of saline (0.9% NaCl) with no Rose Bengal, and followed the same procedure. After injection, mice were prepared for surgery during the 5 minutes allowed for Rose Bengal dye to diffuse. Fur was removed and the scalp was disinfected with alternative swipes of 70% ethanol and betadine. During surgery body temperature was monitored and maintained at 37 ± 0.5°C with a rectal probe and feedback-controlled heating pad. An incision through the midline with surgical scissors was made to expose the skull and locate Bregma. In this study, the brain region to apply photothrombosis is located at 3.28 mm posterior and 2.80 mm lateral to Bregma. After 5 minutes from Rose Bengal injection and location of stereotaxic coordinates, the brain region was illuminated through the skull for 10 minutes with a 520nm diode laser (Thor labs) set at 2mm diameter and 10mW power. Skull was then washed with saline 0.9% NaCl and the incision was closed with Vetbond tissue adhesive (3M 1469Sb). Triple antibiotic and 2% lidocaine were applied to the closed incision, and the mouse was returned to a clean cage.

### Tissue collection and preparation

Mice were injected intraperitoneally with 100 mg/kg Ketamine (Victor Medical Company) and 20 mg/kg Xylazine (Anased) mix and subjected to transcardial perfusion. Perfusion was performed with PBS to obtain fresh frozen tissue for fluorescent *in situ* hybridization (FISH). Those brains were collected, embedded in OCT (Sciegen 4583) and stored at -80°C until analysis. For the rest of experiments, perfusion was performed with PBS followed by 4% PFA (paraformaldehyde) to obtain fixed tissue. Brains were collected and stored in 4% PFA at 4°C overnight, after which they were washed three times with PBS and transferred to 30% sucrose and stored for 3 days at 4°C for cryoprotection. Those brains were embedded in TFM (General data healthcare TFM-5), frozen in dry ice/100% ethanol and stored at -80°C until analysis. For TTC staining, mice were euthanized by intraperitoneal injection of ketamine and xylazine mix and brains were collected without prior perfusion.

### Fluorescent *in situ* hybridization (FISH)

Coronal sections were obtained from WT C57BL/6J mice that went under sham surgeries or photothrombosis and euthanized 24h or 7 days after surgery to analyze expression of Chrdl1. Sections were made at a thickness of 16µm, cut with a cryostat (Hacker Industries OTF5000) at coordinates corresponding with the core of the injury (3.28 mm posterior and 2.80 mm lateral to Bregma). Fluorescent in situ hybridizations (FISH, ACDbio 320850) were performed according to manufacturer’s instructions with some modifications. Brain slices were pre-treated with Protease IV for 20 minutes. Probes used were Chrdl1 (ACDbio 442811), Slc1a3/GLAST (ACDbio 430781-C2). For negative control we used a 3-plex negative probe (ACDbio 31043) to determine the background fluorescence. For mounting, slices were applied with SlowFade gold antifade mountant with DAPI (Thermo Fisher Scientific S36939). On top of the sections we used coverslips 22 mm × 50 mm 1.5 thickness and sealed with nail polish. Slc1a3 was imaged in channel 488, and Chrdl1 in channel 550. Three brain sections were imaged per mouse, and a minimum of 3 mice per condition (sham or stroke). Peri-infarct region in the ipsilateral hemispheres, and the equivalent region in the contralateral hemispheres were imaged on a Zeiss LSM 710 confocal microscope using 20x/0.8NA objective as 16 bit images at 1024 × 1024 pixels (pixel size 0.31×0.31µm) as z-stacks of 5 slices with a total thickness of 6.783 um. Representative images are orthogonal projections. Fluorescence intensity was measured using ImageJ, and calculated as mean intensity of the ROI multiplied by % of area stained.

### 2,3,5-Triphenyltetrazolium chloride (TTC) staining

Chrdl1 KO and WT male mice were subjected to sham surgeries or photothrombosis and brains were extracted without prior perfusion 24h or 7 days later after euthanasia by intraperitoneal injection of overdose of ketamine/xylazine mixture. Brains were cut in 1mm thick slices and incubated in a pre-warm solution of 2% 2,3,5-Triphenyltetrazolium chloride (TTC, Sigma T8877) in PBS for 15 minutes at 37°C, protected from light. Sections were transferred to cold 4% PFA, and later imaged using 0.8X magnification on a stereo zoom microscope (Nikon SMZ-445). Injury and hemisphere volumes were measured with ImageJ, and injury volume was expressed as % of ipsilateral hemisphere. At least 3 biological replicates were used for each time point (24h, 7 days) and condition (sham, stroke), and slices at 0.7, 1.7, 2.7 and 3.7 mm posterior to Bregma were analyzed.

### GFAP and Iba1 staining

YFP-Chrdl1 KO and YFP-WT male mice were subjected to sham surgeries or photothrombosis, and perfused 24h or 7 days later with PBS and 4% PFA to collect fixed tissue. Fixed brains were sliced in the cryostat at a thickness of 16µm and stained for GFAP or Iba1. Upon collection, slices were incubated in blocking solution (5% goat serum, 0.3% Triton X-100 in PBS) for 1h at room temperature. Slices were then incubated with rabbit polyclonal primary antibody anti-GFAP (Abcam ab7260) at a dilution 1:500 or rabbit polyclonal anti-Iba1 at a dilution 1:250 (Wako 016-20001) in antibody buffer (5% goat serum, 100mM lysine, 0.3% Triton X-100 in PBS) overnight at 4°C in a humidified chamber. Slices were washed three times with PBS and incubated with secondary anti-rabbit Alexa 594 (Thermo Fisher Scientific A11012) in antibody buffer at a dilution of 1:500 for 2h at room temperature. Slices were washed three times with PBS and mounted using SlowFade gold antifade mountant with DAPI (Thermo Fisher Scientific S36939), covered with coverslips 22 mm × 50 mm 1.5 thickness and sealed with nail polish. The core and the peri-infarct areas in the ipsilateral hemisphere and equivalent regions in the contralateral hemisphere were imaged using a fluorescent microscope Zeiss Axio Imager.Z2. Images were taken using a 10x/0.45NA objective as 16 bit mosaics of 9 tiles with 10% overlap. To measure fluorescence intensity of GFAP, a region of interest (ROI) of 1000×1000 pixels (pixel size 0.645×0.645µm) was selected in the peri-infarct area, and in the homologous contralateral area (see Figure 1A). For Iba1 fluorescence intensity measurements, an ROI in the core of the injury was selected. Fluorescence intensity was measured using ImageJ, and calculated as mean intensity of the ROI multiplied by % of area stained.

**Figure 1.**
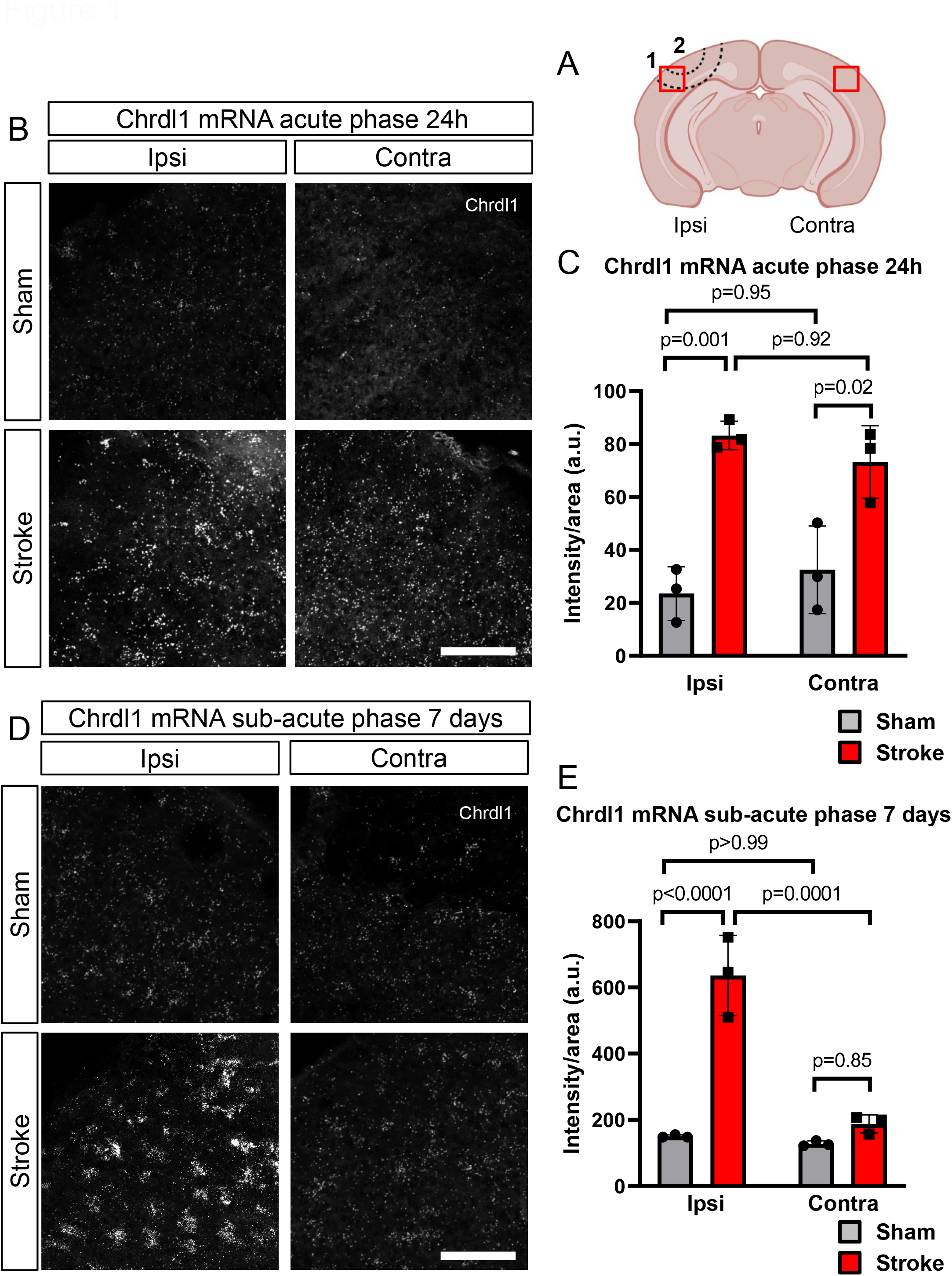
Chrdl1 expression increases in response to ischemic conditions. A) Schematic drawing of a coronal section of a mouse brain showing where the ischemic injury was applied. Region labelled as 1 delimited by two dashed lines indicates the peri-infarct area, and label 2 delimited by a dashed line corresponds to the core, the brain region illuminated with the laser in order to cause the photothrombotic lesion. Red boxes indicate the region of interest (ROI) imaged and analyzed in all experiments in the hemisphere of the stroke (ipsi) and the homologous contralateral region (contra), unless noted differently. B) Representative images of fluorescent *in situ* hybridization (FISH) of Chrdl1 in the peri-infarct area (ipsilateral hemisphere, ipsi) and in the homologous contralateral hemisphere (contra) to the ischemic stroke lesion of coronal sections 24 hours after stroke or sham surgeries. Images were taken in layers 2/3 of the visual cortex. C) Quantification of fluorescence intensity per unit area. Sham mice N=3, stroke N=3. D, E) Same as B, C, 7 days after stroke or sham surgeries. Sham N=3, stroke N=3. Statistics by two-way ANOVA. Scale bar 100µm.

### Spine imaging and analysis

YFP-Chrdl1 KO and YFP-WT male mice were subjected to sham surgeries or photothrombosis and perfused 24h or 7 days later to obtain fixed tissue (see tissue preparation and collection section). Slices were collected from fixed brains in the cryostat at 60µm thickness at coordinates corresponding with the core of the injury (3.28 mm posterior and 2.80 mm lateral to Bregma) and surroundings to be able to image the peri-infarct area. Slices were mounted using SlowFade gold antifade mountant with DAPI (Thermo Fisher Scientific S36939) and coverslips 22 mm × 50 mm 1.5 thickness sealed with nail polish. Peri-infarct region (ipsilateral) and contralateral regions were imaged using a Zeiss LSM 880 Airyscan FAST super-resolution microscope. Images were taken using the 63x/1.4NA oil-immersion objective, as 16 bit images at 1648 × 1648 pixels (pixel size 0.04×0.04µm) as z-stacks of 30 steps with a total thickness of 5.51µm. Representative images are orthogonal projections. Images were analyzed using NeuronStudio software (Computational Neurobiology and Imaging Center CNIC, Mount Sinai School of Medicine, NY). Measurements were from secondary dendrites from neurons on layer 5 extending into layers 2/3 of the visual cortex in the more ventral region of the peri-infarct area and the equivalent region in the contralateral hemisphere. At least 6 dendrites per mouse were analyzed. Spine classifier within the software was set as 1.1µm head-to-neck ratio threshold, 2.5µm height-to-width ratio threshold and mushroom head size of 0.35µm or larger in order to classify spines as immature (stubby or thin), or mature (mushroom) (23).

### Quantification and statistical analysis

Mice were randomly assigned to sham or stroke groups, and the experimenter was blind to genotype at the time of analysis. Sample size for each experiment was based on previous studies in the literature. No animals were excluded from the study.

GraphPad Prism 8 (San Diego, CA) was used to design graphs and statistical analysis. *In situ* hybridizations in Figure 1 and Sup Fig 1; TTC staining in Figure 2; immunostaining experiments in Figure 3, 4 and Sup Fig 3 and 4; and spine density in Figure 5B,F and Sup Figure 5B,F were analyzed using 2-way ANOVA with Sidak’s post-hoc test. One-way ANOVA with Tukey’s multiple comparison test was used to analyze the spine morphology dataset in Figure 5C,D,G,H and Sup Figure 5C,D,G,H. Results are shown as mean ± S.D. with dots representing mice, and graphs show exact p-values. Full statistical calculations are presented in Sup Table 1.

**Figure 2.**
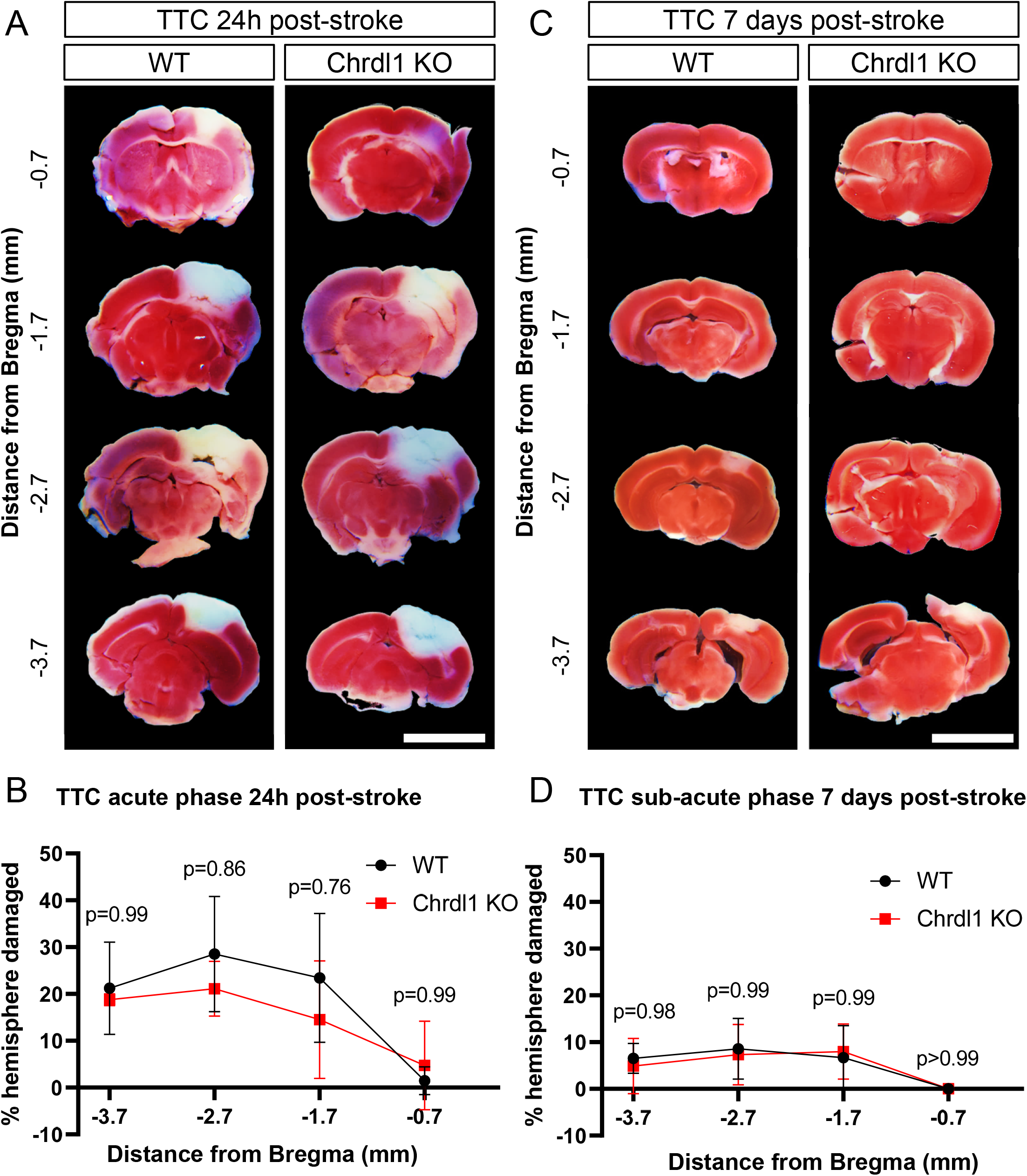
Absence of Chrdl1 does not affect injury volume. A) Representative images of 1mm-thick coronal sections of WT and Chrdl1 KO mouse brains 24 hours after stroke stained with TTC, corresponding to 0.7, 1.7, 2.7 and 3.7mm posterior from Bregma. The dead tissue (brain regions infarcted) is not stained by TTC and appears white, whereas the rest of the tissue turns a shade of red. B) Quantification of the volume of the injury as percentage of the total volume of the ipsilateral hemisphere. WT N=4, Chrdl1 KO N=4. C, D) Same as A, B, 7 days after stroke stained with TTC. WT N=4, Chrdl1 KO N=6. Statistics by two-way repeated measures ANOVA. Scale bar 5mm.

**Figure 3.**
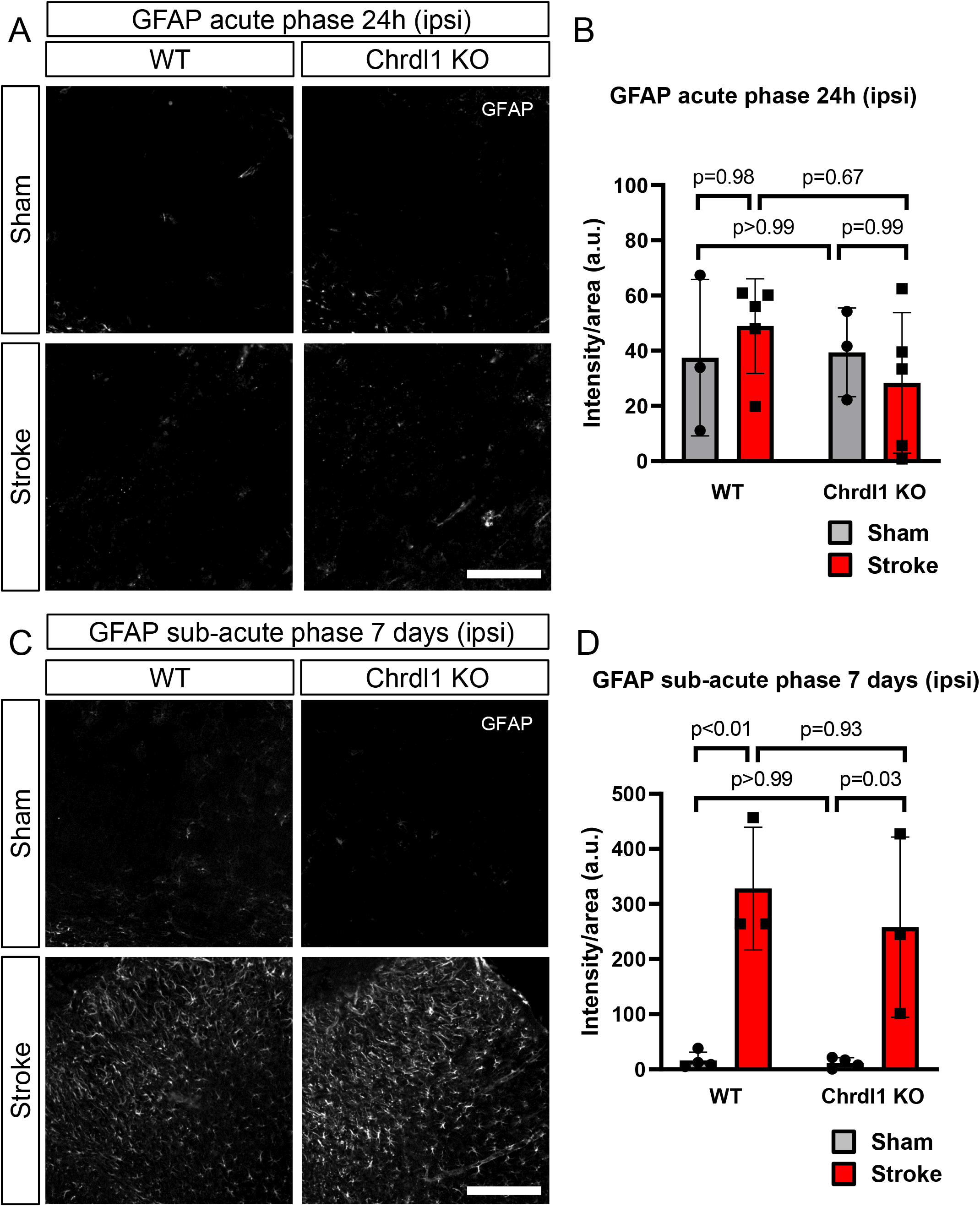
Reactive astrogliosis is not impaired in Chrdl1 KO mice after stroke. A) Representative images of the peri-infarct area of WT and Chrdl1 KO mouse cortex 24 hours after stroke or sham surgery immunostained for GFAP. B) Quantification of GFAP intensity per area unit. WT sham N=3, WT stroke N=5, Chrdl1 KO sham N=3 and Chrdl1 KO stroke N=5. C, D) Same as A, B, 7 days after stroke or sham surgery immunostained for GFAP. WT sham N=4, WT stroke N=3, Chrdl1 KO sham N=4 and Chrdl1 KO stroke N=3. Statistics by two-way ANOVA. Scale bar 200µm.

**Figure 4.**
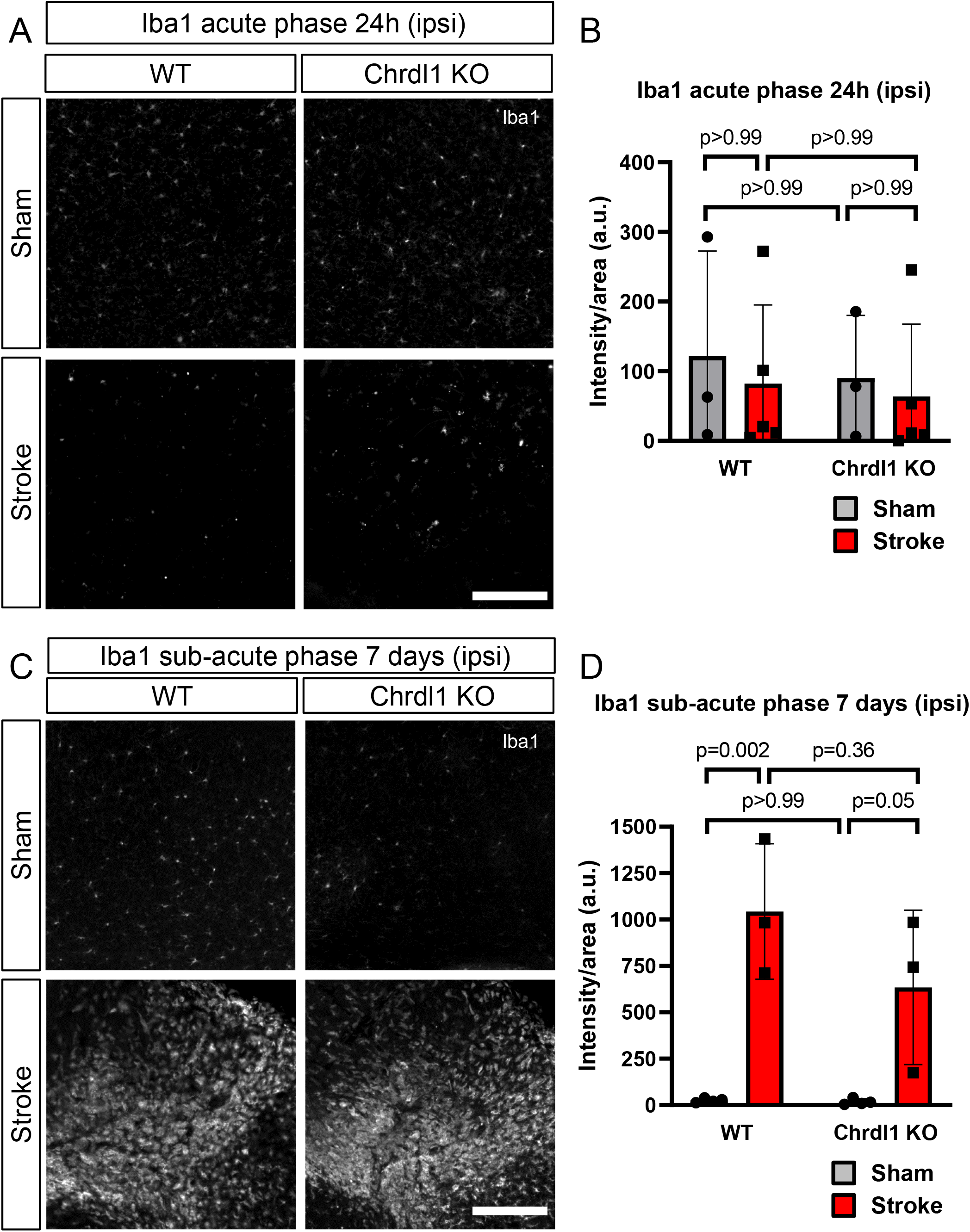
Absence of Chrdl1 does not affect the microglia response after stroke. A) Example images of the injury core of WT and Chrdl1 KO mouse cortex 24 hours after stroke or sham surgery immunostained for Iba1. B) Quantification of Iba1 intensity per area unit. WT sham N=3, WT stroke N=5, Chrdl1 KO sham N=3 and Chrdl1 KO stroke N=5. C, D) Same as A, B, 7 days after stroke or sham surgery immunostained for Iba1. WT sham N=4, WT stroke N=3, Chrdl1 KO sham N=4 and Chrdl1 KO stroke N=3. Statistics by two-way ANOVA. Scale bar 200µm.

**Figure 5.**
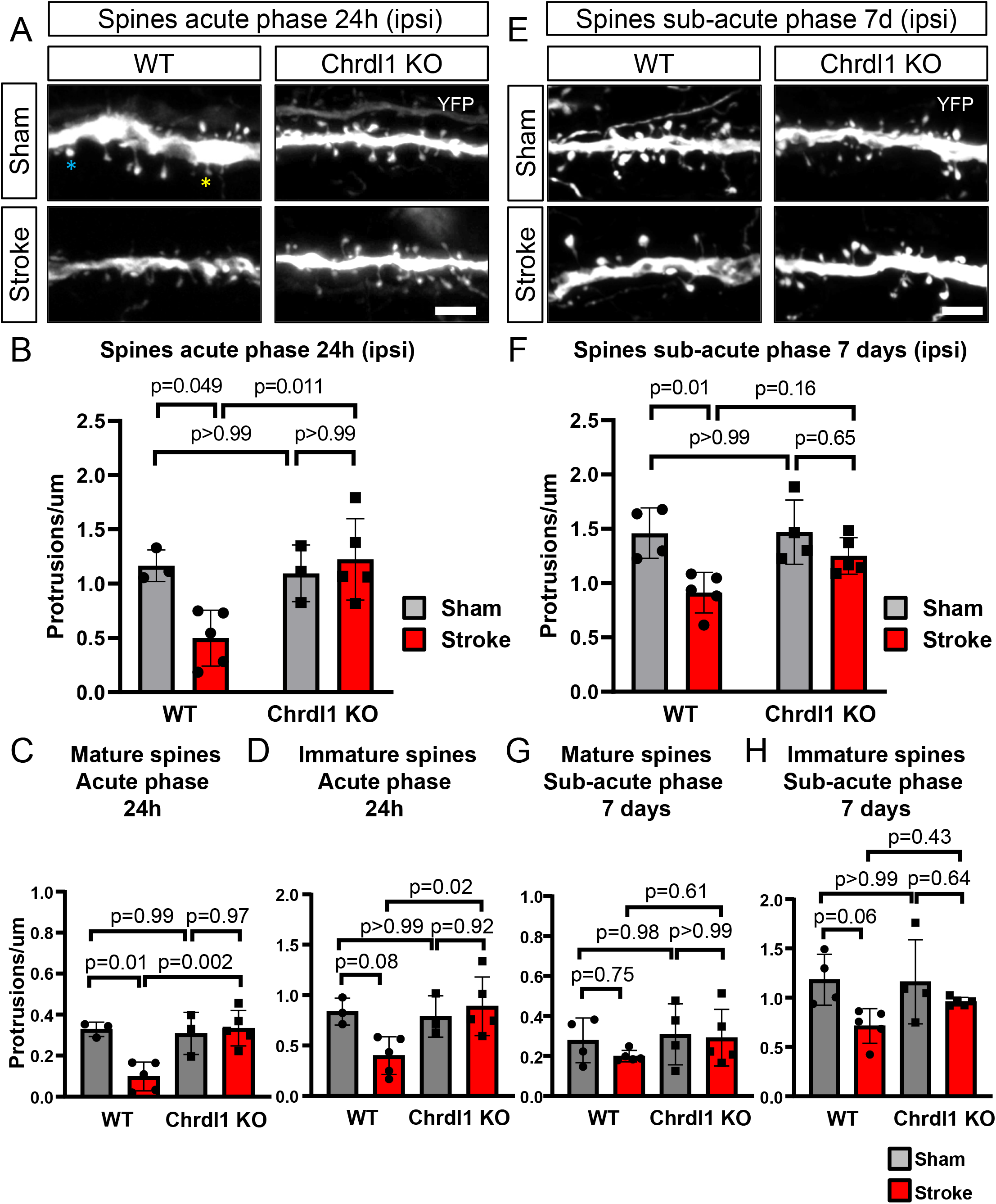
Absence of Chrdl1 prevents spine loss in the peri-infarct area. A) Representative images of YFP expressing layer 5 neuron secondary dendrites present in layers 2/3 of the visual cortex in WT or Chrdl1 KO mice, 24 hours after stroke or sham surgeries. The blue star indicates an example of a spine of mature morphology, and the yellow star indicates an example of a spine of immature morphology. B) Quantification of number of spines per µm in dendrites in layers 2/3 in the peri-infarct area of the visual cortex of mice 24 hours after stroke or sham surgery. WT sham N=3, WT stroke N=5, Chrdl1 KO sham N=3, Chrdl1 KO stroke N=5. Statistics by two-way ANOVA. C,D) Quantification of spines with mature (C) or immature (D) morphology in the peri-infarct area of dendrites in layers 2/3 of the visual cortex of mice 24 hours after stroke or sham surgery. WT sham N=3, WT stroke N=5, Chrdl1 KO sham N=3, Chrdl1 KO stroke N=5. Statistics by one-way ANOVA. E) Same as A, 7 days after stroke or sham surgery. F) Same as B, 7 days after stroke or sham surgery. WT sham N=4, WT stroke N=5, Chrdl1 KO sham N=4, Chrdl1 KO stroke N=5. Statistics by two-way ANOVA. G,H) Same as C, D, 7 days after stroke or sham surgery. WT sham N=4, WT stroke N=5, Chrdl1 KO sham N=4, Chrdl1 KO stroke N=5. Statistics by one-way ANOVA. Scale bars 5µm.

## Results

### Chrdl1 expression increases in response to ischemic conditions

In response to ischemic stroke caused by MCAO (middle cerebral artery occlusion), the expression of various astrocytic genes is altered (24). SPARC and TSP-1, both astrocyte-secreted factors with synaptogenic roles, are upregulated during the acute phase and they have been linked to functional recovery (17, 25). Here we used photothrombotic stroke to cause ischemia in the visual cortex of 4 month old male mice to study the role of Chrdl1. The effects were analyzed during the acute (24 hours post-stroke) and sub-acute (7 days post-stroke) phases, two stages post-injury characterized by different plasticity levels. The sub-acute phase is considered the time window during which, given the enhanced endogenous plasticity, recovery and neural repair is facilitated (6).

Due to its role in synaptic plasticity, and previous microarray data showing that Chrdl1 expression is increased by MCAO (24), we hypothesized that Chrdl1 expression could be regulated in response to ischemic stroke in confined areas of the injured brain. By fluorescence *in situ* hybridization (FISH), we analyzed RNA expression of Chrdl1 in the peri-infarct area (ipsilateral, ipsi) and in the homologous contralateral (contra) hemisphere (Figure 1A). We found that expression of Chrdl1, which we previously demonstrated to be astrocyte-specific (18), was upregulated in astrocytes in the peri-infarct area of WT mice during the acute phase (24 hours after insult) (Figure 1B,C, ipsi: sham 23.5 ± 5.8 a.u., stroke, 83.2 ± 3.1 a.u.). This increase in the expression of Chrdl1 was also observed in non-injured tissue on the contralateral hemisphere to the lesion, (Figure 1B,C, contra: sham, 32.5 ± 9.6 a.u., stroke, 73.2 ± 7.9 a.u.). During the sub-acute phase (7 days post-stroke) the expression of Chrdl1 in the contralateral hemisphere was restored to physiological levels (Figure 1D,E contra: sham, 127.9 ±4.8 a.u., stroke, 187.7 ± 15.6 a.u.), whereas the increased expression in the peri-infarct area was persistent and significantly increased (Figure 1D,E ipsi: sham, 150.1 ± 2.6 a.u., stroke, 636.3 ± 70.1 a.u.). This suggests that the upregulation of astrocytic Chrdl1 after ischemic stroke may trigger mechanisms that hinder synaptic plasticity, obstructing synaptic regeneration and hampering recovery.

We also analyzed the expression of the astrocyte-enriched gene Slc1a3 (Glutamate Aspartate Transporter, GLAST) to see whether the changes in expression that we observed in Chrdl1 were specific to Chrld1 or shared by other astrocyte genes. GLAST was chosen due to its important roles in glutamate uptake in the context of excitotoxic injuries such as ischemic stroke (26). We found that GLAST was stable in the peri-infarct area (Sup Figure 1A,B ipsi: sham, 362.4 ± 52.9 a.u., stroke, 445.2 ± 21.5 a.u.) and in the homologous contralateral region during the acute phase (Sup Figure 1A,B, contra: sham 319.2 ± 45.3 a.u., stroke 375.5 ± 83.9 a.u.). During the sub-acute phase, and similar to what we observed with the expression of Chrdl1, GLAST expression increased in the peri-infarct area (Sup Figure 1C,D, ipsi: sham, 561.8 ± 35.7 a.u., stroke, 913.5 ± 119.5 a.u.), while in the homologous contralateral region there was no significant upregulation (Sup Figure 1C,D contra: sham, 552.2 ± 31.9 a.u., stroke, 855.4 ± 56.1 a.u.). These results demonstrate that both Chrdl1 and GLAST expression are upregulated in response to stroke but with distinct time courses, with Chrdl1 upregulation showing more rapid changes than GLAST in response to the same sort of injury.

### Absence of Chrdl1 does not affect injury volume

One factor that determines the severity of the injury is the size of the brain region affected by the deprivation of blood supply. Increasing plasticity can help repair and remap neuronal circuits that were damaged during the ischemic episode (27). It has been shown that manipulating astrocytic or neuronal plasticity-regulating molecules contributes to reducing the size of the injury and improved functional recovery (17, 20). In our previous study we found that constitutive Chrdl1 KO mice display enhanced experience-dependent plasticity in the visual system (18), thus we hypothesized that Chrdl1 KO mice would show decreased injury volume after an ischemic lesion. To address this we performed photothrombosis in male Chrdl1 KO (-/y) and WT (+/y) mice. However, absence of Chrdl1 did not affect the volume of the injury during the acute phase (Figure 2A,B, 24h post-stroke, WT: −0.7mm from Bregma 1.5 ± 1.5%, -1.7mm from Bregma 23.4 ± 6.9%, -2.7mm from Bregma 28.5 ± 12.3%, -4.7mm from Bregma 21.2 ± 4.9%, Chrdl1 KO: −0.7mm from Bregma 4.7 ± 4.7%, -1.7mm from Bregma 14.5 ± 6.3%, -2.7mm from Bregma 21.1 ± 2.9%, -4.7mm from Bregma 18.8 ± 2.6%). The injury volume during the sub-acute phase was also unaltered in the absence of Chrdl1 (Figure 2C,D, 7days post-stroke, WT: −0.7mm from Bregma 0.0 ± 0.0%, -1.7mm from Bregma 6.7 ± 3.4%, - 2.7mm from Bregma 8.6 ± 3.3%, -4.7mm from Bregma 6.5 ± 1.6%, Chrdl1 KO: −0.7mm from Bregma 0.0 ± 0.0%, -1.7mm from Bregma 8.0 ± 2.4%, -2.7mm from Bregma 7.3 ± 2.6%, - 4.7mm from Bregma 4.9 ± 2.4%). As expected, mice subjected to sham surgeries did not show any injury (Sup Figure 2). This demonstrates that Chrdl1 does not impact the progression of the size of the core of the injury, and any potential role for Chrdl1 may reside in the surrounding tissue corresponding to the peri-infarct area.

### Reactive astrogliosis is not impaired in Chrdl1 KO mice after stroke

As a consequence of ischemic stroke, astrocytes become reactive in the peri-infarct area and undergo molecular, morphological and functional changes that drive the formation of the glial scar surrounding the core of the injury (28). GFAP (glial fibrillary acidic protein) is a cytoskeletal protein that is upregulated in astrocytes in response to different types of CNS injury, and it is an indicator of reactive astrogliosis (28). We measured GFAP intensity to assess whether Chrdl1 played a role in attenuation or intensification of reactive astrogliosis in the peri-infarct area of WT and Chrdl1 KO mice. During the acute phase we did not detect significant increases in GFAP in either WT or Chrdl1 KO peri-infarct areas (Figure 3A,B, 24h post-stroke, WT: sham, 37.5 ± 14.2 a.u., stroke, 48.9 ± 9.9 a.u., Chrdl1 KO: sham, 39.4 ± 8.1 a.u., stroke, 28.3 ± 14.7 a.u.). During the sub-acute phase we detected an increased intensity in GFAP staining which was significant, but independent of Chrdl1 expression (Figure 3C,D, 7 days post-stroke, WT: sham, 16.0 ± 7.6 a.u., stroke, 328.1 ± 64.1 a.u., Chrdl1 KO: sham, 11.7 ± 4.6 a.u., stroke, 257.9 ± 94.3 a.u.). There were no variations in GFAP levels in the contralateral hemisphere during the acute phase (Sup Figure 3A,B 24h post-stroke, WT: sham, 33.1 ± 11.6 a.u., stroke, 24.8 ± 16.2 a.u., Chrdl1 KO: sham, 44.5 ± 14.6 a.u., stroke, 14.9 ± 10.8 a.u.) or sub-acute phase (Sup Figure 3C,D 7days post-stroke, WT: sham, 5.3 ± 1.6 a.u., stroke, 7.5 ± 1.2 a.u., Chrdl1 KO: sham, 16.6 ± 7.8 a.u., stroke, 8.7 ± 3.7 a.u.). Our results indicate that Chrdl1 does not regulate GFAP, a protein involved in reactive astrogliosis and in the glial scar formation in certain pathological contexts (29), suggesting that Chrdl1 may not play relevant roles in reactive astrogliosis.

### Absence of Chrdl1 does not affect the microglia response after stroke

Microglia have important roles in neuroinflammation, however, it is unclear what determines their beneficial or detrimental function in response to ischemic stroke. In response to stroke, Iba1+ microglial cells increase in the core of the injury which indicates activation of a microglial response (30). We assessed if there was a role for Chrdl1 in microglia activation by performing Iba1 staining of WT and Chrdl1 KO tissue after stroke. In the core of the injury Iba1 fluorescent signal was not evident during the first 24 hours post-stroke corresponding to the acute phase (Figure 4A,B, 24h post-stroke, WT: sham, 121.5 ± 75.4 a.u., stroke, 82.0 ± 65.4 a.u., Chrdl1 KO: sham, 90.0 ± 45.0 a.u., stroke, 63.8 ± 59.9 a.u.), but it was apparent at 7 days post-stroke, during the sub-acute phase, in both WT and Chrdl1 KO mice (Figure 4C,D, 7days post-stroke, WT: sham, 25.4 ± 5.6 a.u., stroke, 1043.3 ± 210.8 a.u., Chrdl1 KO: sham, 17.5 ± 8.2 a.u., stroke, 634.3 ± 240.1 a.u.). In both WT and Chrdl1 KO mice, differences in microglia activation were not observed in the contralateral hemisphere during the acute phase (Sup Figure 4A,B 24h post-stroke, WT: sham, 115.7 ± 65.8 a.u., stroke, 76.1 ± 45.9 a.u., Chrdl1 KO: sham, 94.0 ± 46.8 a.u., stroke, 108.2 ± 79.4 a.u.) or the sub-acute phase (Sup Figure 4C,D 7days post-stroke, WT: sham, 27.1 ± 7.5 a.u., stroke, 30.5 ± 15 a.u., Chrdl1 KO: sham, 14.2 ± 5.0 a.u., stroke, 20.6 ± 8.4 a.u.). Our results indicate that Chrdl1 does not have a large effect on the microglia response to ischemic stroke.

### Absence of Chrdl1 prevents spine loss in the peri-infarct area

Immediately after ischemic stroke, there is an acute loss of dendritic spines in the peri-infarct area in WT mice. The decreased spine density is severe during the acute phase, and persistent at later stages with signs of potential recovery during the sub-acute phase (22). However, the exact mechanisms that promote recovery of spine density are not completely understood. The absence of Chrdl1 does not affect spine density or morphology in uninjured adult mice (18), but we hypothesized that it may have an effect in response to ischemic stroke, as was observed in other studies where plasticity-regulating proteins were eliminated (31). To this end, WT and Chrdl1 KO mice where layer 5 neurons express the fluorescent protein YFP were subjected to sham surgeries or photothrombotic stroke in the visual cortex, and spine density of secondary dendrites extending into layers 2/3 of the visual cortex were analyzed. WT mice subjected to photothrombotic stroke displayed severe loss of spines in the peri-infarct area during the acute phase (Figure 5A,B, 24h post-stroke, WT: sham, 1.17 ± 0.08 protrusion/µm, stroke, 0.50 ± 0.11 protrusion/µm), a well-documented consequence of ischemic stroke (22, 32, 33). Interestingly, Chrdl1 KO mice did not show the characteristic ischemic-induced decrease in spine density during the acute phase (Figure 5A,B, 24h post-stroke, Chrdl1 KO: sham, 1.10 ± 0.15 protrusion/µm, stroke, 1.22 ± 0.17 protrusion/µm), suggesting that absence of Chrdl1 prevents the initial degeneration of synaptic structures in the peri-infarct area. We observed that spine loss in the peri-infarct area was persistent in WT mice during the sub-acute phase, though not as severe as initially observed (Figure 5E,F 7 days post-stroke, WT: sham, 1.46 ± 0.12 protrusion/µm, stroke, 0.91 ± 0.08 protrusion/µm). During the sub-acute phase, Chrdl1 KO mice displayed spine numbers similar to those mice that went under sham surgeries (Figure 5E,F 7 days post-stroke, Chrdl1 KO: sham, 1.47 ± 0.15 protrusion/µm, stroke, 1.25 ± 0.08 protrusion/µm), suggesting that absence of Chrdl1 protects from spine loss rather than delaying the neurodegenerative processes that drive spine elimination after stroke. Neither WT nor Chrdl1 KO mice showed variation in spine density in the contralateral hemisphere (Sup Figure 5A,B,E,F).

One possibility is that Chrdl1 KO mice experience spine loss with a fast turnover during the early acute phase (first 24 hours post-insult), in which case an increase in immature spine morphologies would be expected compared to mice that went under sham surgeries. We found that WT mice showed a significant decrease in mature spines after stroke (Figure 5C, WT: sham 0.33 ± 0.02 protrusion/µm, stroke, 0.10 ± 0.03 protrusion/µm). Chrdl1 KO did not show any alterations in mature spine density in the peri-infarct area during the acute phase (Figure 5C, Chrdl1 KO: sham, 0.31 ± 0.06 protrusion/µm, stroke, 0.33 ± 0.04 protrusion/µm). During the acute phase, WT mice also showed a decrease in immature spine density (Figure 5D, WT: sham 0.84 ± 0.08 protrusion/µm, stroke, 0.40 ±0.08 protrusion/µm), whereas Chrdl1 KO mice had a constant density of immature spines after stroke compared to sham animals (Figure 5D, Chrdl1 KO: sham, 0.79 ± 0.12 protrusion/µm, stroke, 0.89 ± 0.13 protrusion/µm). During the sub-acute phase, the WT mice showed restoration of mature spine density (Figure 5G, WT: sham, 0.28 ± 0.06 protrusion/µm, stroke, 0.20 ± 0.01 protrusion/µm), while the Chrdl1 KO mice showed no changes in the mature spine density (Figure 5G, Chrdl1 KO: sham, 0.31 ± 0.08 protrusion/µm, stroke, 0.29 ± 0.06 protrusion/µm). On the other hand, density of immature spines in the WT mice during the sub-acute phase remained low (Figure 5H, WT: sham 1.18 ± 0.13 protrusion/µm, stroke, 0.71 ± 0.08 protrusion/µm). During the sub-acute phase Chrdl1 KO mice did not experience delayed loss of immature spines as the density of the spines with immature morphology remained at comparable levels to the acute phase (Figure 5H, Chrdl1 KO: sham, 1.16 ± 0.21 protrusion/µm, stroke, 0.96 ± 0.02 protrusion/µm). There were no significant morphological changes in the contralateral hemisphere of WT or Chrdl1 KO in response to stroke (Sup Figure 5C,D,G,H). This indicates that the absence of Chrdl1 protected the peri-infarct neurons from the typical spine loss observed in WT mice in response to ischemic stroke, rather than promoting more rapid recovery of spines.

## Discussion

In this study we investigated the contribution of the astrocyte-enriched synapse-regulating protein Chrdl1 to protection from ischemic stroke. We found that Chrdl1 is upregulated after stroke, and that removal of Chrdl1 prevents neuronal dendritic spine loss that is characteristic of the acute phase after injury. We further found that this protective role of Chrdl1 is independent of reactive gliosis and does not impact on the initial size of the injury.

### Astrocytic Chrdl1 expression increases in response to ischemic stroke

We found that Chrdl1 expression is upregulated in the acute phase after stroke in the peri-infarct area and in the contralateral hemisphere, and that the upregulation is persistent during the sub-acute phase only in the peri-infarct area (Figure 1B-E). Contralateral alterations are possible in response to a focal injury as observed before (34) but the mechanisms are yet to be fully described. We also found that the glutamate aspartate transporter (GLAST) is increased in expression in the sub-acute phase, but not the acute phase. In ischemia there is a massive release of glutamate into the synaptic cleft that ultimately triggers excitotoxicity and cellular death, and astrocytes play important roles in re-uptake of excessive glutamate including through GLAST (35). The differential temporal changes of Chrdl1 and GLAST expression in response to stroke suggest that dysregulation of these factors may be persistent at later timepoints during the chronic phase (30+ days post-stroke) and that other astrocyte factors may also display differential temporal regulation in response to the same kind of injury. The increased expression of Chrdl1 may limit plasticity during the sub-acute phase, a time when post-stroke endogenous plasticity is enhanced (5), as suggested by the apparent stability of dendritic spines observed in Chrdl1 KO mice. In the future it will be important to assess astrocyte gene expression during the acute, sub-acute and chronic phases, as they may dictate the endogenous plasticity levels that determine functional recovery and circuit remodeling in response to injury.

### Progression of injury volume is independent of Chrdl1 expression

The size of the injury has important implications in the functional outcome from ischemic stroke. Normally, larger injury sizes are more likely to have fatal outcomes (36). Blockade of Chrdl1 promotes enhanced experience-dependent plasticity in the brains of mice (18) and so we hypothesized that absence of Chrdl1 may facilitate recovery from stroke through reduction of injury volume, as has been shown for other plasticity limiting molecules (20). We found, however, that absence of Chrdl1 did not have any effects on the volume of the injury (Figure 2, Sup Figure 2). This is in contrast to a study where Noggin, a BMP (Bone Morphogenetic Protein) antagonist similar to Chrdl1, was over-expressed and shown to promote reduction in injury size and improved behavioral outcomes (37), suggesting differences in mode of action of these different BMP inhibitors. Interestingly, a previous study on thrombospondin-1 and -2, two astrocyte-secreted synaptogenic proteins, showed that their absence did not play a role in the size of the injury, but impaired functional recovery (17). In the future it will be interesting to assess the levels of other proteins that have been previously identified to regulate the size of the injury during ischemic stroke, and assess whether their expression is altered in Chrdl1 KO mice to trigger mechanisms that maintain injury volume.

### Chrdl1 does not impact reactive gliosis

Glial cells respond to diverse types of injury and disease through a series of molecular and morphological changes that group under the term reactive gliosis. In the case of astrocytes, reactive astrogliosis in its most severe form leads to the formation of the glial scar (28). According to our results based on GFAP immunohistochemistry, Chrdl1 absence does not have an effect on reactive astrogliosis (Figure 3, Sup Figure 3), suggesting that scar formation is Chrdl1-independent in the context of ischemic stroke. This can explain why the absence of Chrdl1 does not affect the volume of the injury in Chrdl1 KO mice when compared to WT mice (Figure 2), as the formation of the glial scar is not affected by knocking out Chrdl1. The effects of Chrdl1 appear to be related to neuronal plasticity mechanisms in response to external stimuli, such as sensory deprivation (18) or injury, rather than intrinsic mechanisms of astrocytes themselves that lead to reactive astrogliosis and formation of the glial scar. Further experiments to analyze other reactive markers like Lcn2 or Serpina3n among others (24) will clarify whether Chrdl1 regulates astrogliosis in ischemic stroke.

The immune response in ischemic stroke is regulated by various glial cell types that also interact among themselves to regulate inflammation, and these glia-mediated mechanisms can exert both beneficial and detrimental effects (13). Astrocytes and microglia are two important components of the neurovascular unit, and both are activated in response to ischemic stroke, as well as other types of CNS injuries (38). It has been previously shown that activated microglia can induce reactive astrogliosis with neurotoxic effects (39), but the mechanisms that regulate microglia activation are still not completely clear. In the present study we found that absence of Chrdl1 does not affect microglia activation, as demonstrated by Iba1 immunostaining (Figure 4, Sup Figure 4). This suggests that Chrdl1 does not play a role in the inflammatory response in stroke, an important feature that leads to further damage beyond the core and the peri-infarct areas, contributing to the extension of the injury and worse outcomes. It has been previously reported that other BMP antagonists, such as Noggin, play important roles in the microglia response in ischemic stroke (37). As Chrdl1 is also a BMP antagonist it will be important to determine if it regulates microglia activation at later time points post-stroke for example during the chronic phase, when typically the microglial response is reduced. Therefore, the beneficial effects of Chrdl1 removal on dendritic spines do not appear to be related to inflammation or astrogliosis.

### Absence of Chrdl1 prevents spine loss in the peri-infarct area after ischemic stroke

In response to ischemic stroke neurons in the peri-infarct region undergo deep remodeling, featuring spine loss and morphological modifications which are especially severe during the acute phase (22). During later stages post-stroke, spontaneous mechanisms promote spinogenesis, but those mechanisms remain unclear. Since we found an increased expression of Chrdl1 in response to ischemic injury that may hinder synaptic remodeling, we hypothesized that eliminating Chrdl1 could protect spine loss or promote faster spine turnover to compensate for the ischemic damage. We found that Chrdl1 KO mice did not show the characteristic spine loss in the peri-infarct area in response to ischemic stroke during either acute or sub-acute phases (Figure 5, Sup Figure 5). We did not observe any alteration in spine morphology, suggesting that absence of Chrdl1 protects spines from ischemic-driven pruning rather than promoting a quick turnover in response to ischemic lesion. In the future live imaging experiments will help to determine whether spines in Chrdl1 KO mice are stable and protected in response to ischemic stroke. Therefore time- and region-dependent Chrdl1 upregulation may hinder remodeling of damaged circuits by preventing formation and/or restructuring of neuronal synapses.

### Chrdl1 may play different roles depending on the post-stroke phase

Our work shows that increased expression of Chrdl1 can be a detrimental contribution from astrocytes that impedes recovery by limiting plasticity and circuit remodeling. But how can Chrdl1 hinder recovery? As outlined above, one possibility is by blocking BMP signaling mechanisms. It has been previously shown that administration of BMP7 24 hours after ischemic stroke improves behavioral recovery (40). Post-stroke upregulation of Chrdl1, a known BMP antagonist with affinity for BMP4, 6 and 7 (41), could be blocking the beneficial effects of BMPs after ischemic stroke. Blockade of Chrdl1 after ischemic stroke may therefore represent a new approach to promote the neurorestorative roles of BMPs. However, different BMPs may play diverse roles in stroke, promoting recovery or delayed cell death. In a different study it was found that over-expressing the BMP antagonist Noggin had neuroprotective effects by reducing the size of the injury (37). Noggin has high affinity for BMP4, like Chrdl1 (41). Therefore it is important to determine the specific post-stroke phase in which to manipulate Chrld1 in order to promote beneficial effects.

An alternative explanation for the protective effect of Chrdll1 after stroke is through its role as a regulator of synapse maturation and synaptic AMPAR composition (18). Excessive glutamate release in response to ischemic stroke stimulates GluA2-lacking AMPARs, which are permeable to Ca^2+^, a process known as excitotoxicity that triggers injurious signals that ultimately lead to delayed neuronal death (42). This occurs immediately following ischemic injury, and later effects on GluA2-containing AMPARs have not been thoroughly analyzed. The presence of the subunit GluA2 renders the AMPAR impermeable to Ca^2+^, and thus upregulation of Chrdl1 may play a beneficial role following excitotoxic injury (i.e. during the acute phase post-stroke) as it increases the number of Ca^2+^-impermeable GluA2-containing AMPARs on the neuronal surface (18). However, high expression of Chrdl1 at later stages post-stroke (for example during the sub-acute phase as we observed) could lead to limited plasticity, preventing synaptic remodeling and ultimately recovery. These previous studies and our own results suggest that Chrdl1 may be playing different roles at the various stages post-stroke and have variable effects on different neuronal populations, and so in the future it will be important to analyze the expression of Chrdl1 at later time points during chronic (30+ days post-stroke) phase post-stroke and in other brain regions.

In conclusion, our results indicate that astrocyte-secreted factors play a role in circuit remodeling after ischemic insult, and Chrdl1 in particular may hamper recovery during the sub-acute phase due to its role in limiting plasticity in the cortex. The restricted regional expression of Chrdl1 suggests that alternative astrocyte-secreted factors that regulate plasticity and spine density in other brain regions may also mediate plasticity mechanisms in response to region-specific ischemic injury in a time-dependent manner. This study highlights the importance of dissecting astrocyte-mediated plasticity mechanisms to understand the limitations in circuit remodeling in response to injury and other pathologies of the CNS, and design new strategies to promote neuroprotection and recovery at different time points post-stroke.

## Acknowledgements

We thank Cari Dowling for technical assistance, Dr. Amy Gleichman and Dr. S. Thomas Carmichael for assistance with the photothrombotic technique, and Dr. Uri Manor for advice with imaging acquisition. This work was supported by NIH NINDS grant NS105742 to NJA. Work in the lab of N.J.A. is supported by the Hearst Foundation, the Pew Foundation, and the CZI Neurodegeneration Network. This work was supported by Core Facilities of the Salk Institute (Biophotonics: NIH NCI CCSG P30 014195, the Waitt, Helmsley and Chapman Foundations).

## Author contribution statement

EB-S designed and performed experiments and conducted analysis, with input from NJA. EB-S and NJA conceived the project and wrote the manuscript.

## Supplementary Information

Supplementary material consisting of 5 Figures and 1 Table can be found at the journal website.

