## Supplementary Figures 1-5 for "Astrocyte-secreted chordin-like 1 regulates spine density after ischemic stroke"

Supplemental figure 1, related to main figure 1

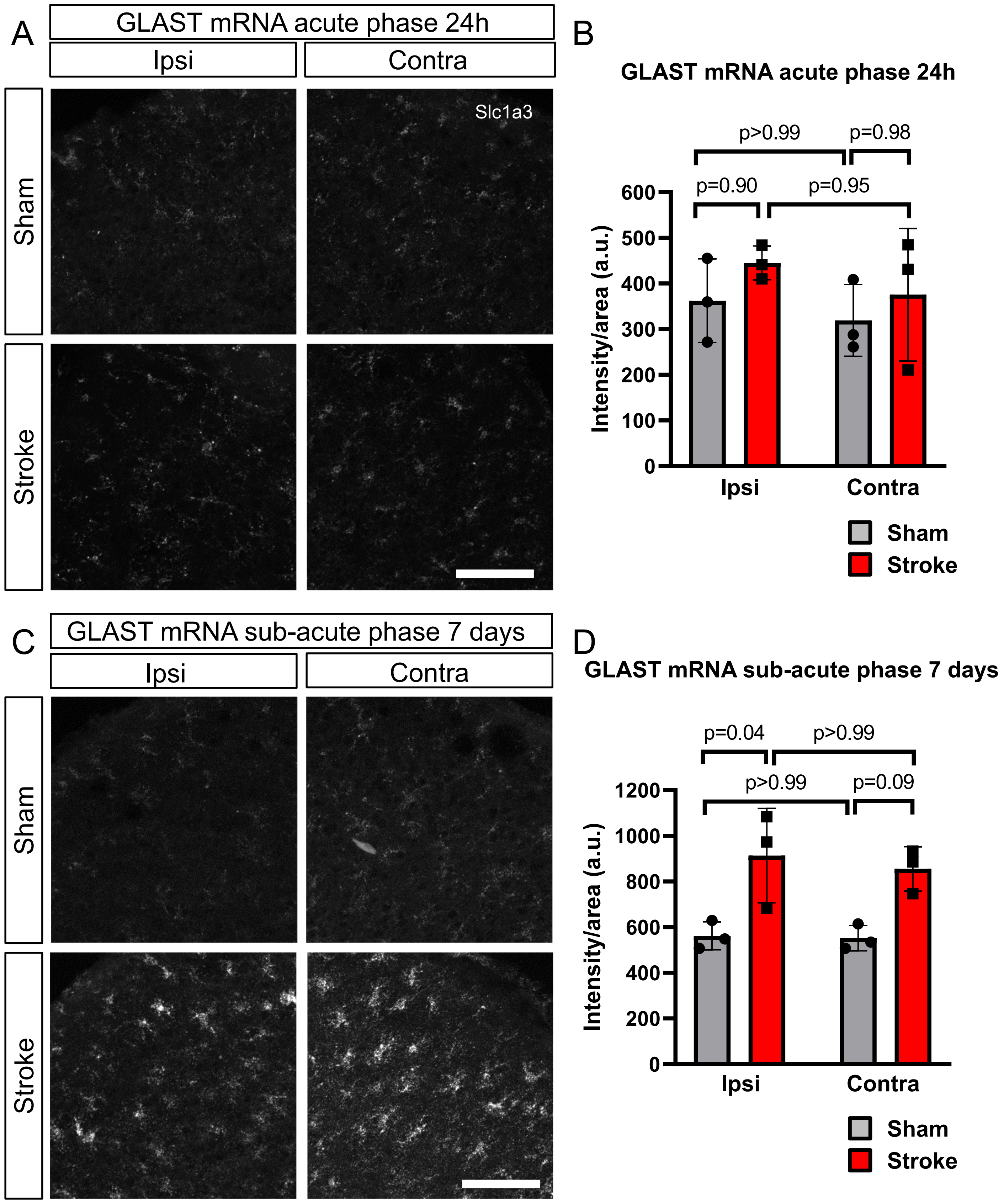

**Supplemental figure 1 (related to main figure 1) GLAST expression increases in the peri—infarct area only during sub-acute phase.** A) Representative images of FISH of GLAST (Slc1a3) in the peri-infarct area (ipsilateral hemisphere, ipsi) and in the contralateral hemisphere (contra) to the ischemic stroke lesion of coronal sections 24 hours after stroke or sham surgeries. Images from layers 2/3 of the visual cortex. B) Quantification of fluorescence intensity per unit area. Sham mice N=3, stroke N=3. C, D) Same as A, B, 7 days after stroke or sham surgeries. Sham N=3, stroke N=3. Statistics by two-way ANOVA. Scale bar 100μm.

Supplemental figure 2, related to main figure 2

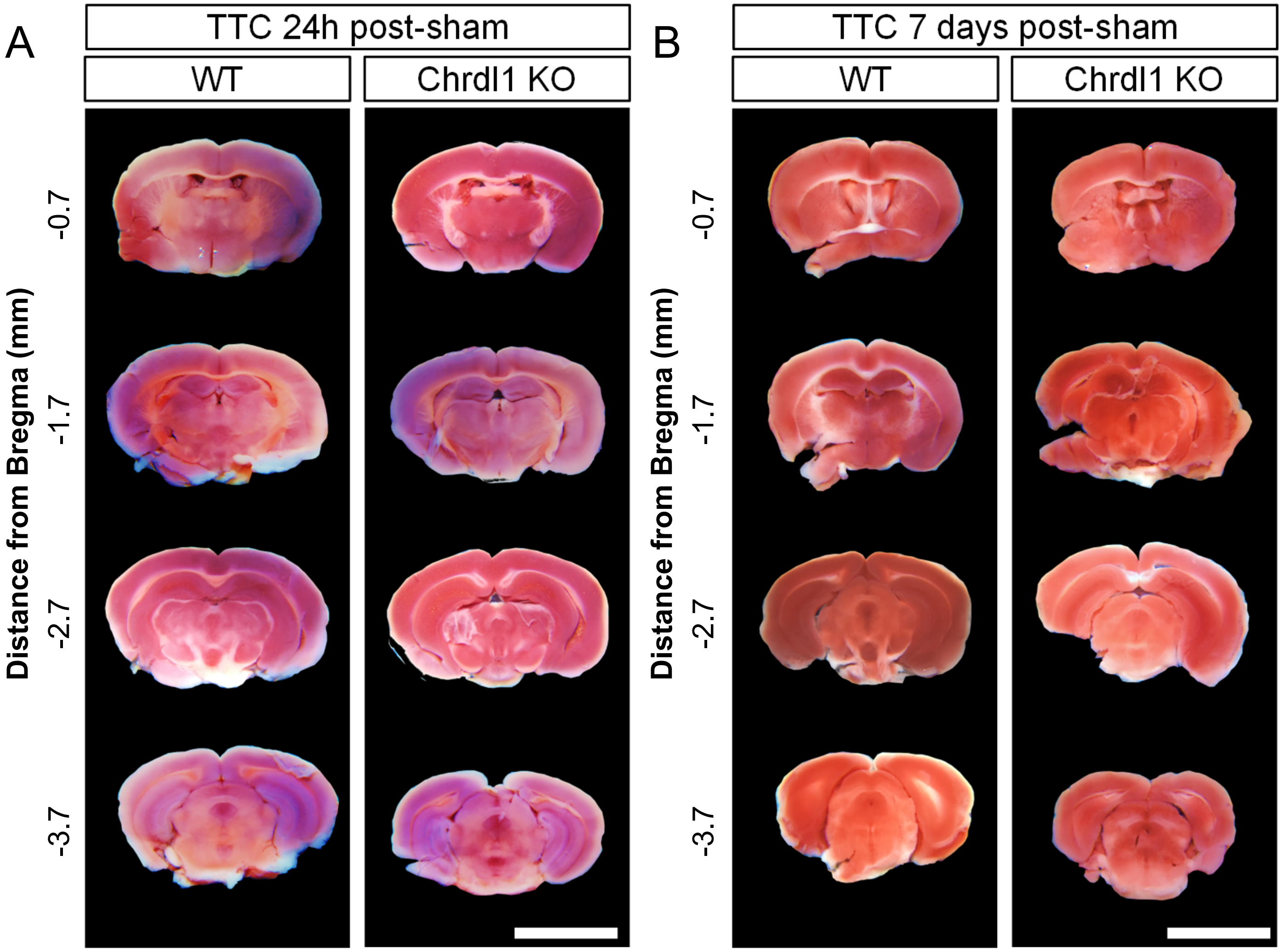


**Supplemental figure 2 (related to main figure 2) Absence of Chrdl1 does not affect injury volume.** A) Representative images of 1mm-thick coronal sections of WT and Chrdl1 KO mouse brains 24 hours after sham surgeries. Slices correspond to 0.7, 1.7, 2.7 and 3.7mm posterior from Bregma as indicated. Brain slices were stained with TTC. No infarct tissue is present. WT sham N=4, Chrdl1 KO N= 3 B) Representative images of the same experiment but 7 days after sham surgeries. WT sham N=4, Chrdl1 KO N=4. Scale bar 5mm.

Supplemental figure 3, related to main figure 3

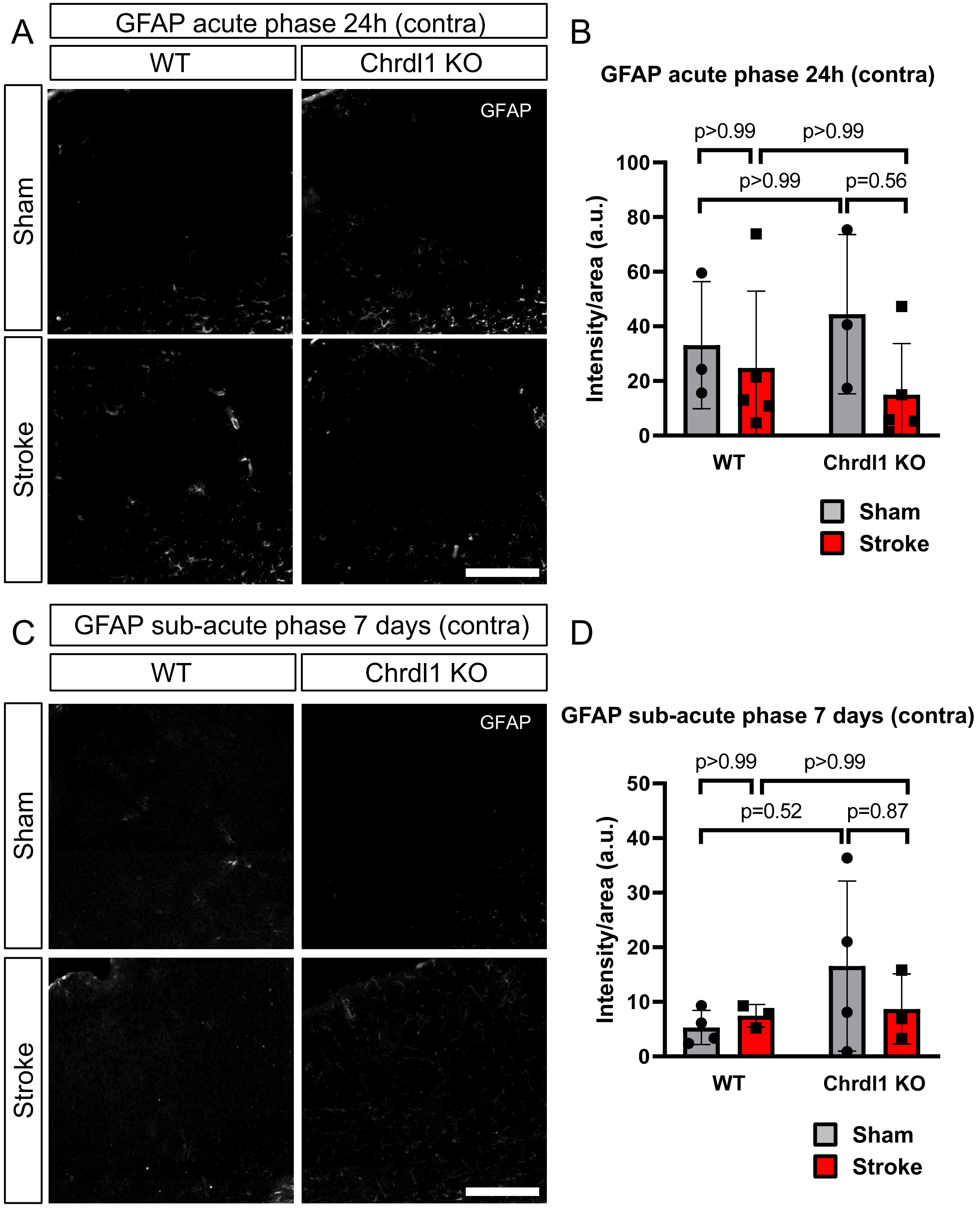

**Supplemental figure 3 (related to main figure 3) Reactive astrogliosis is not impaired in Chrdl1 KO mice after stroke.** A) Representative images of the ROI in the homologous contralateral hemisphere to the stroke of WT and Chrdl1 KO mice 24 hours after stroke or sham surgeries immunostained for GFAP B) Quantification of GFAP intensity per area unit. WT sham N=3, WT stroke N=5, Chrdl1 KO sham N=3 and Chrdl1 KO stroke N=5. C, D) Same as A, B 7 days after stroke or sham surgeries. WT sham N=4, WT stroke N=3, Chrdl1 KO sham N=4 and Chrdl1 KO stroke N=3. Statistics by two-way ANOVA. Scale bar 200μm.

Supplemental figure 4, related to main figure 4

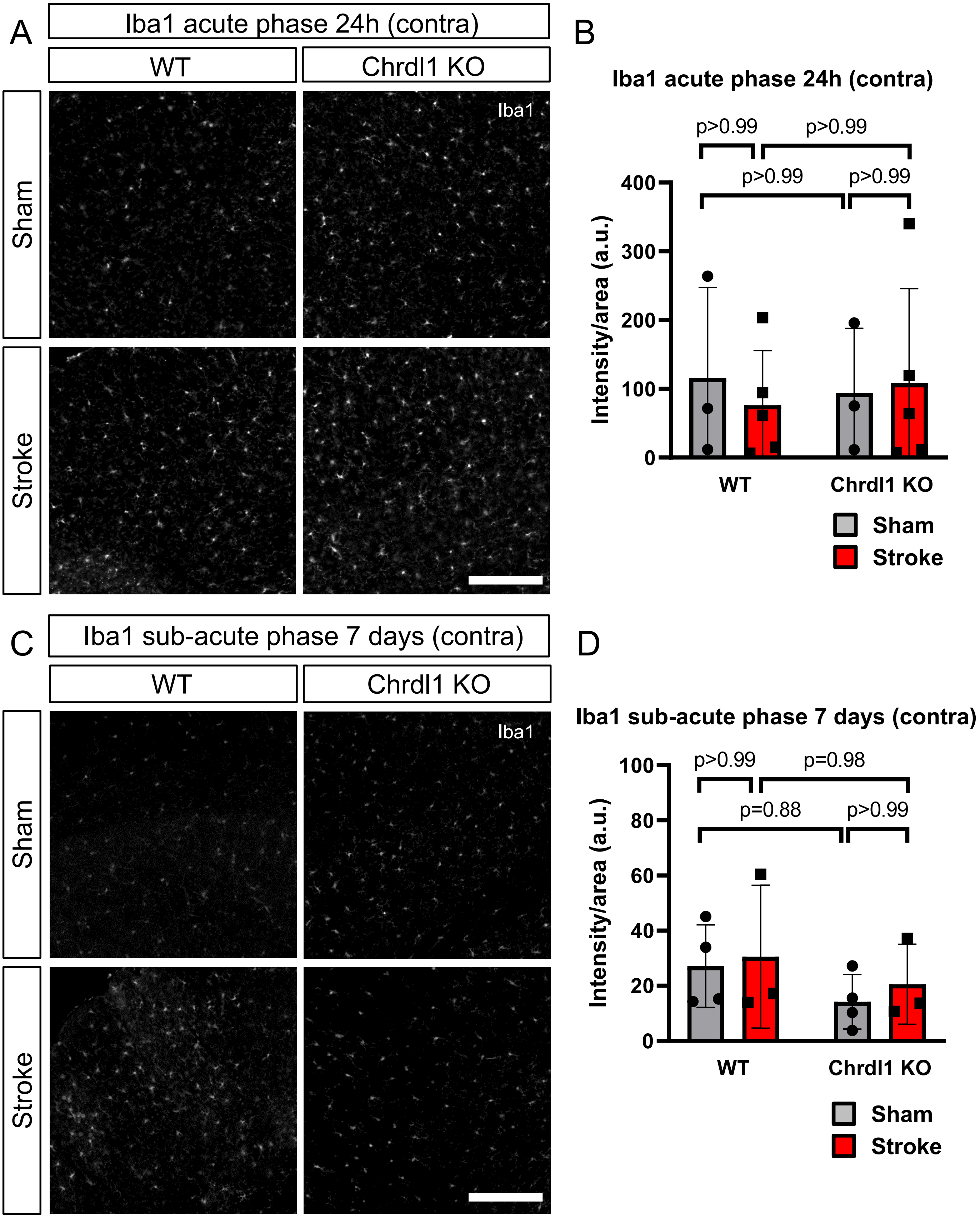

**Supplemental figure 4 (related to main figure 4) Absence of Chrdl1 does not affect microglia response after stroke.** A) Representative images showing Iba1 immunostaining of the ROI in the homologous contralateral hemisphere to the stroke of WT and Chrdl1 KO mice 24 hours after stroke or sham surgeries. B) Quantification of Iba1 intensity per area unit. WT sham N=3, WT stroke N=5, Chrdl1 KO sham N=3 and Chrdl1 KO stroke N=5. C, D) Same as A, B, 7 days after stroke or sham surgeries. WT sham N=4, WT stroke N=3, Chrdl1 KO sham N=4 and Chrdl1 KO stroke N=3. Statistics by two-way ANOVA. Scale bar 200µm.

Supplemental figure 5, related to main figure 5

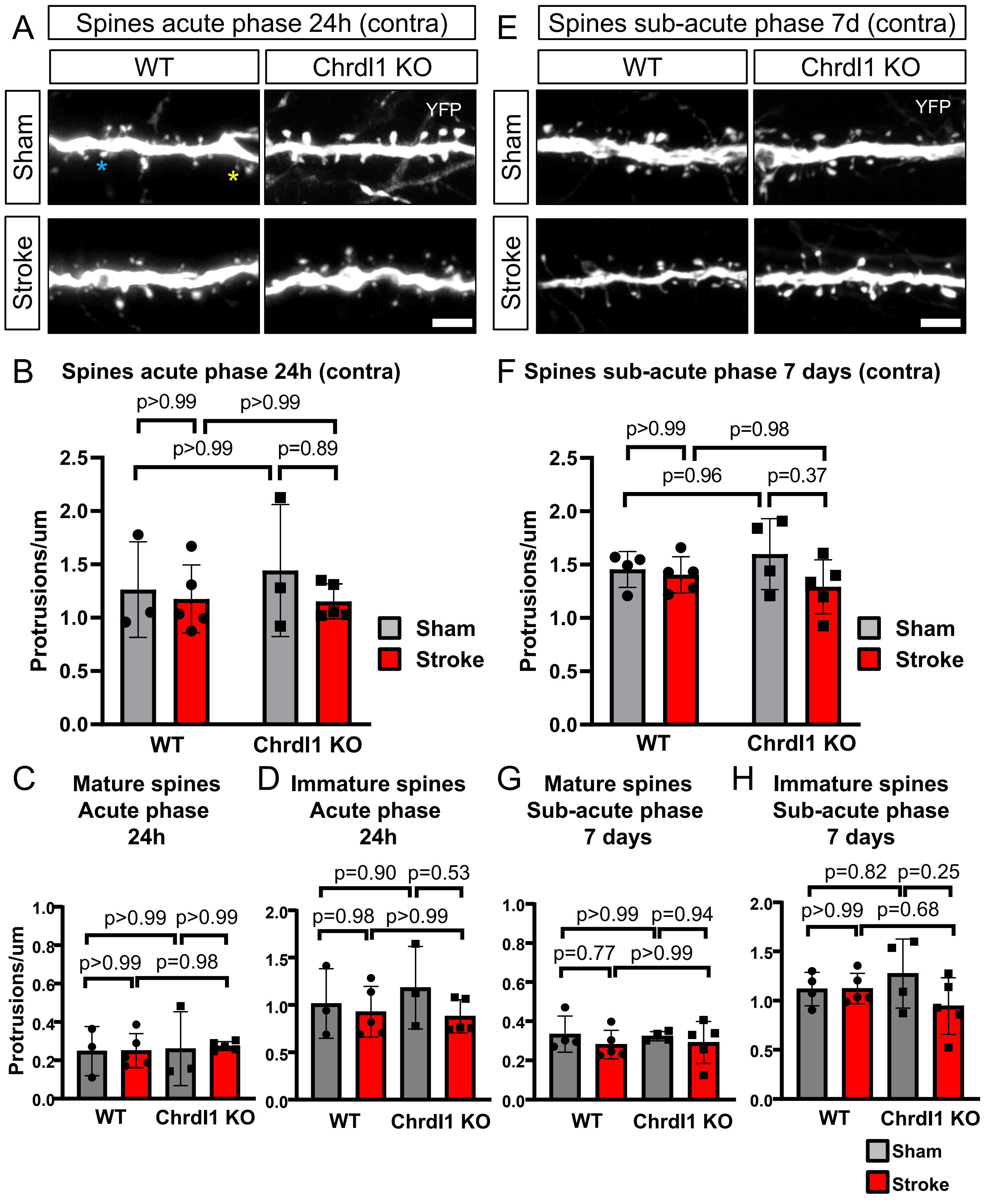

**Supplemental figure 5 (related to main figure 5) Absence of Chrdl1 prevents spine loss in the peri-infarct area.** A) Representative images of YFP expressing layer 5 neuron secondary dendrites present in layers 2/3 of the visual cortex in WT or Chrdl1 KO mice, 24 hours after stroke or sham surgeries in the contralateral side to the stroke. Blue star indicates an example of a spine of mature morphology, and the yellow star indicates an example of a spine of immature morphology. B) Quantification in the contralateral area of number of spines per μm in dendrites in layers 2/3 of the visual cortex of animals 24 hours after stroke or sham surgery. WT sham N=3, WT stroke N=5, Chrdl1 KO sham N=3, Chrdl1 KO stroke N=5. Statistics by two-way ANOVA. C,D) Quantification in the contralateral area of spines with mature (C) or immature (D) morphology on dendrites in layers 2/3 of the visual cortex of mice 24 hours after stroke or sham surgery. WT sham N=3, WT stroke N=5, Chrdl1 KO sham N=3, Chrdl1 KO stroke N=5. Statistics by one-way ANOVA. E) Same as A, 7 days after stroke or sham surgeries in the contralateral area to the stroke. F) Same as B, 7 days after stroke or sham surgeries in the contralateral area to the stroke. WT sham N=4, WT stroke N=5, Chrdl1 KO sham N=4, Chrdl1 KO stroke N=5. Statistics by two-way ANOVA. G,H) Same as C, D, 7 days after stroke or sham surgery. WT sham N=4, WT stroke N=5, Chrdl1 KO sham N=4, Chrdl1 KO stroke N=5. Statistics by one-way ANOVA. Scale bars 5μm.
