## Supplemental Table 1 for "Astrocyte-secreted chordin-like 1 regulates spine density after ischemic stroke"

**Chrdl1 intensity x %area** Figure 1

**24 hours post-stroke**

| IPSI | Mean | Std. Error of Mean | N | Šídák's multiple comparisons test | Predicted (LS) mean diff. | 95.00% CI of diff. | Below threshold? | Summary | Adjusted P Value |
|---|---|---|---|---|---|---|---|---|---|
| WT sham | 23.5 | 5.8 | 3 | ipsi:Sham vs. ipsi:Stroke | -59.73 | -94.14 to -25.32 | Yes | ** | 0.0019 |
| WT stroke | 83.2 | 3.1 | 3 | ipsi:Sham vs. contra:Sham | -8.98 | -43.39 to 25.43 | No | ns | 0.9497 |
| CONTRA | | | | ipsi:Sham vs. contra:Stroke | -49.72 | -84.13 to -15.31 | Yes | ** | 0.0063 |
| WT sham | 32.5 | 9.6 | 3 | ipsi:Stroke vs. contra:Sham | 50.75 | 16.34 to 85.16 | Yes | ** | 0.0055 |
| WT stroke | 73.2 | 7.9 | 3 | ipsi:Stroke vs. contra:Stroke | 10.00 | -24.41 to 44.41 | No | ns | 0.9199 |
| | | | | contra:Sham vs. contra:Stroke | -40.74 | -75.15 to -6.333 | Yes | * | 0.0204 |

**7 days post-stroke**

| IPSI | Mean | Std. Error of Mean | N | Šídák's multiple comparisons test | Predicted (LS) mean diff. | 95.00% CI of diff. | Below threshold? | Summary | Adjusted P Value |
|---|---|---|---|---|---|---|---|---|---|
| WT sham | 150.2 | 2.6 | 3 | ipsi:Sham vs. ipsi:Stroke | -486.10 | -662.7 to -309.6 | Yes | **** | <0.0001 |
| WT stroke | 636.3 | 70.1 | 3 | ipsi:Sham vs. contra:Sham | 22.22 | -154.3 to 198.8 | No | ns | 0.9988 |
| CONTRA | | | | ipsi:Sham vs. contra:Stroke | -37.50 | -214.0 to 139.1 | No | ns | 0.9809 |
| WT sham | 128.0 | 4.8 | 3 | ipsi:Stroke vs. contra:Sham | 508.30 | 331.8 to 684.9 | Yes | **** | <0.0001 |
| WT stroke | 187.7 | 15.6 | 3 | ipsi:Stroke vs. contra:Stroke | 448.60 | 272.1 to 625.2 | Yes | *** | 0.0001 |
| | | | | contra:Sham vs. contra:Stroke | -59.72 | -236.3 to 116.8 | No | ns | 0.8547 |

**GLAST intensity x %area** Supplementary figure 1

**24 hours post-stroke**

| IPSI | Mean | Std. Error of Mean | N | Šídák's multiple comparisons test | Predicted (LS) mean diff. | 95.00% CI of diff. | Below threshold? | Summary | Adjusted P Value |
|---|---|---|---|---|---|---|---|---|---|
| WT sham | 362.4 | 53.0 | 3 | ipsi:Sham vs. ipsi:Stroke | -82.85 | -355.2 to 189.5 | No | ns | 0.9035 |
| WT stroke | 445.2 | 21.5 | 3 | ipsi:Sham vs. contra:Sham | 43.21 | -229.1 to 315.5 | No | ns | 0.9957 |
| CONTRA | | | | ipsi:Sham vs. contra:Stroke | -13.15 | -285.5 to 259.2 | No | ns | >0.9999 |
| WT sham | 319.2 | 45.3 | 3 | ipsi:Stroke vs. contra:Sham | 126.10 | -146.3 to 398.4 | No | ns | 0.6161 |
| WT stroke | 375.5 | 83.9 | 3 | ipsi:Stroke vs. contra:Stroke | 69.70 | -202.6 to 342.0 | No | ns | 0.9539 |
| | | | | contra:Sham vs. contra:Stroke | -56.36 | -328.7 to 216.0 | No | ns | 0.9832 |

**7 days post-stroke**

| IPSI | Mean | Std. Error of Mean | N | Šídák's multiple comparisons test | Predicted (LS) mean diff. | 95.00% CI of diff. | Below threshold? | Summary | Adjusted P Value |
|---|---|---|---|---|---|---|---|---|---|
| WT sham | 561.8 | 35.7 | 3 | ipsi:Sham vs. ipsi:Stroke | -351.70 | -695.7 to -7.703 | Yes | * | 0.0447 |
| WT stroke | 913.5 | 119.5 | 3 | ipsi:Sham vs. contra:Sham | 9.67 | -334.3 to 353.7 | No | ns | >0.9999 |
| CONTRA | | | | ipsi:Sham vs. contra:Stroke | -293.50 | -637.5 to 50.47 | No | ns | 0.1046 |
| WT sham | 552.2 | 31.9 | 3 | ipsi:Stroke vs. contra:Sham | 361.40 | 17.37 to 705.3 | Yes | * | 0.0389 |
| WT stroke | 855.4 | 56.1 | 3 | ipsi:Stroke vs. contra:Stroke | 58.17 | -285.8 to 402.2 | No | ns | 0.9940 |
| | | | | contra:Sham vs. contra:Stroke | -303.20 | -647.2 to 40.80 | No | ns | 0.0908 |

**TTC staining** Figure 2

**24 hours post-stroke**

WT — **%hemisphere damage**

| **Distance from Bregma** | **Mean** | **Std. Error of Mean** | **N** |
|---|---|---|---|
| **-0.7** | 1.5 | 1.5 | 4 |
| **-1.7** | 23.4 | 6.9 | 4 |
| **-2.7** | 28.5 | 6.2 | 4 |
| **-3.7** | 21.2 | 4.9 | 4 |

Chrdl1 KO — **%hemisphere damage**

| **Distance from Bregma** | **Mean** | **Std. Error of Mean** | **N** |
|---|---|---|---|
| **-0.7** | 4.7 | 4.7 | 4 |
| **-1.7** | 14.5 | 6.3 | 4 |
| **-2.7** | 21.1 | 2.9 | 4 |
| **-3.7** | 18.8 | 0.6 | 4 |

**Šídák's multiple comparisons test**

WT - Chrdl1 KO

| **Mean Diff.** | **95.00% CI of diff.** | **Below threshold?** | **Summary** | **Adjusted P Value** |
|---|---|---|---|---|
| -3.22 | -27.38 to 20.93 | No | ns | 0.9922 |
| 8.92 | -15.23 to 33.08 | No | ns | 0.7617 |
| 7.39 | -16.76 to 31.55 | No | ns | 0.8601 |
| 2.44 | -21.72 to 26.59 | No | ns | 0.9973 |

**7 days post-stroke**

WT — **%hemisphere damage**

| **Distance from Bregma** | **Mean** | **Std. Error of Mean** | **N** |
|---|---|---|---|
| **-0.7** | 0.0 | 0.0 | 4 |
| **-1.7** | 6.7 | 3.4 | 3 |
| **-2.7** | 8.6 | 3.3 | 4 |
| **-3.7** | 6.5 | 1.6 | 4 |

Chrdl1 KO — **%hemisphere damage**

| **Distance from Bregma** | **Mean** | **Std. Error of Mean** | **N** |
|---|---|---|---|
| **-0.7** | 0.0 | 0.0 | 6 |
| **-1.7** | 8.0 | 2.4 | 5 |
| **-2.7** | 7.3 | 2.6 | 6 |
| **-3.7** | 4.9 | 2.4 | 5 |

**Šídák's multiple comparisons test**

WT - Chrdl1 KO

| **Predicted (LS) mean diff.** | **95.00% CI of diff.** | **Below threshold?** | **Summary** | **Adjusted P Value** |
|---|---|---|---|---|
| -1.78E-15 | -8.659 to 8.659 | No | ns | >0.9999 |
| -1.31 | -11.11 to 8.485 | No | ns | 0.9943 |
| 1.23 | -7.424 to 9.893 | No | ns | 0.9927 |
| 1.68 | -7.320 to 10.68 | No | ns | 0.9801 |

**GFAP intensity x %area (Ipsi)** Figure 3

**24 hours post-stroke**

| IPSI | Mean | Std. Error of Mean | N |
|---|---|---|---|
| WT sham | 37.5 | 14.2 | 3 |
| WT stroke | 48.9 | 9.9 | 5 |
| Chrdl1 KO sham | 39.4 | 8.1 | 3 |
| Chrdl1 KO stroke | 28.3 | 14.7 | 5 |

| Šídák's multiple comparisons test | Predicted (LS) mean diff. | 95.00% CI of diff. | Below threshold? | Summary | Adjusted P Value |
|---|---|---|---|---|---|
| WT:Sham vs. WT:Stroke | -11.46 | -62.24 to 39.33 | No | ns | 0.9828 |
| WT:Sham vs. Chrdl1 KO:Sham | -1.90 | -58.69 to 54.88 | No | ns | >0.9999 |
| WT:Sham vs. Chrdl1 KO:Stroke | 9.12 | -41.67 to 59.90 | No | ns | 0.9948 |
| WT:Stroke vs. Chrdl1 KO:Sham | 9.55 | -41.24 to 60.34 | No | ns | 0.9933 |
| WT:Stroke vs. Chrdl1 KO:Stroke | 20.57 | -23.41 to 64.56 | No | ns | 0.6671 |
| Chrdl1 KO:Sham vs. Chrdl1 KO:Stroke | 11.02 | -39.77 to 61.81 | No | ns | 0.9859 |

**7 days post-stroke**

| IPSI | Mean | Std. Error of Mean | N |
|---|---|---|---|
| WT sham | 16.0 | 7.6 | 4 |
| WT stroke | 328.1 | 64.1 | 3 |
| Chrdl1 KO sham | 11.7 | 4.6 | 4 |
| Chrdl1 KO stroke | 257.9 | 94.3 | 3 |

| Šídák's multiple comparisons test | Predicted (LS) mean diff. | 95.00% CI of diff. | Below threshold? | Summary | Adjusted P Value |
|---|---|---|---|---|---|
| WT:Sham vs. WT:Stroke | -312.10 | -533.6 to -90.52 | Yes | ** | 0.0059 |
| WT:Sham vs. Chrdl1 KO:Sham | 4.34 | -200.8 to 209.5 | No | ns | >0.9999 |
| WT:Sham vs. Chrdl1 KO:Stroke | -241.90 | -463.4 to -20.31 | Yes | * | 0.0305 |
| WT:Stroke vs. Chrdl1 KO:Sham | 316.40 | 94.86 to 538.0 | Yes | ** | 0.0053 |
| WT:Stroke vs. Chrdl1 KO:Stroke | 70.21 | -166.6 to 307.1 | No | ns | 0.9287 |
| Chrdl1 KO:Sham vs. Chrdl1 KO:Stroke | -246.20 | -467.8 to -24.65 | Yes | * | 0.0275 |

**GFAP intensity x %area (Contra)** Supplementary figure 3

**24 hours post-stroke**

| CONTRA | Mean | Std. Error of Mean | N |
|---|---|---|---|
| WT sham | 33.1 | 11.6 | 3 |
| WT stroke | 24.8 | 16.2 | 5 |
| Chrdl1 KO sham | 44.5 | 14.6 | 3 |
| Chrdl1 KO stroke | 14.9 | 10.8 | 5 |

| Šídák's multiple comparisons test | Predicted (LS) mean diff. | 95.00% CI of diff. | Below threshold? | Summary | Adjusted P Value |
|---|---|---|---|---|---|
| WT:Sham vs. WT:Stroke | 8.35 | -48.41 to 65.11 | No | ns | 0.9982 |
| WT:Sham vs. Chrdl1 KO:Sham | -11.34 | -74.80 to 52.11 | No | ns | 0.9949 |
| WT:Sham vs. Chrdl1 KO:Stroke | 18.19 | -38.57 to 74.94 | No | ns | 0.9127 |
| WT:Stroke vs. Chrdl1 KO:Sham | -19.69 | -76.45 to 37.07 | No | ns | 0.8795 |
| WT:Stroke vs. Chrdl1 KO:Stroke | 9.84 | -39.31 to 58.99 | No | ns | 0.9907 |
| Chrdl1 KO:Sham vs. Chrdl1 KO:Stroke | 29.53 | -27.23 to 86.29 | No | ns | 0.5607 |

**7 days post-stroke**

| CONTRA | Mean | Std. Error of Mean | N |
|---|---|---|---|
| WT sham | 5.3 | 1.6 | 4 |
| WT stroke | 7.5 | 1.2 | 3 |
| Chrdl1 KO sham | 16.6 | 7.8 | 4 |
| Chrdl1 KO stroke | 8.7 | 3.7 | 3 |

| Šídák's multiple comparisons test | Predicted (LS) mean diff. | 95.00% CI of diff. | Below threshold? | Summary | Adjusted P Value |
|---|---|---|---|---|---|
| WT:Sham vs. WT:Stroke | -2.16 | -25.13 to 20.81 | No | ns | 0.9998 |
| WT:Sham vs. Chrdl1 KO:Sham | -11.28 | -32.54 to 9.990 | No | ns | 0.5168 |
| WT:Sham vs. Chrdl1 KO:Stroke | -3.39 | -26.36 to 19.58 | No | ns | 0.9978 |
| WT:Stroke vs. Chrdl1 KO:Sham | -9.12 | -32.09 to 13.85 | No | ns | 0.7818 |
| WT:Stroke vs. Chrdl1 KO:Stroke | -1.23 | -25.79 to 23.33 | No | ns | >0.9999 |
| Chrdl1 KO:Sham vs. Chrdl1 KO:Stroke | 7.89 | -15.08 to 30.86 | No | ns | 0.8702 |

**Iba1 intensity x %area (Ipsi)** Figure 4

**24 hours post-stroke**

| IPSI | Mean | Std. Error of Mean | N |
|---|---|---|---|
| WT sham | 121.5 | 75.4 | 3 |
| WT stroke | 82.0 | 65.4 | 5 |
| Chrdl1 KO sham | 90.0 | 45.0 | 3 |
| Chrdl1 KO stroke | 63.8 | 59.9 | 5 |

| Šídák's multiple comparisons test | Predicted (LS) mean diff. | 95.00% CI of diff. | Below threshold? | Summary | Adjusted P Value |
|---|---|---|---|---|---|
| WT:Sham vs. WT:Stroke | 39.47 | -222.1 to 301.1 | No | ns | 0.9980 |
| WT:Sham vs. Chrdl1 KO:Sham | 31.45 | -261.0 to 323.9 | No | ns | 0.9997 |
| WT:Sham vs. Chrdl1 KO:Stroke | 57.72 | -203.9 to 319.3 | No | ns | 0.9846 |
| WT:Stroke vs. Chrdl1 KO:Sham | -8.02 | -269.6 to 253.6 | No | ns | >0.9999 |
| WT:Stroke vs. Chrdl1 KO:Stroke | 18.25 | -208.3 to 244.8 | No | ns | >0.9999 |
| Chrdl1 KO:Sham vs. Chrdl1 KO:Stroke | 26.27 | -235.3 to 287.9 | No | ns | 0.9998 |

**7 days post-stroke**

| IPSI | Mean | Std. Error of Mean | N |
|---|---|---|---|
| WT sham | 25.4 | 5.6 | 4 |
| WT stroke | 1043.3 | 210.8 | 3 |
| Chrdl1 KO sham | 17.5 | 8.2 | 4 |
| Chrdl1 KO stroke | 634.3 | 240.1 | 3 |

| Šídák's multiple comparisons test | Predicted (LS) mean diff. | 95.00% CI of diff. | Below threshold? | Summary | Adjusted P Value |
|---|---|---|---|---|---|
| WT:Sham vs. WT:Stroke | -1018.00 | -1636 to -400.3 | Yes | ** | 0.0019 |
| WT:Sham vs. Chrdl1 KO:Sham | 7.82 | -564.0 to 579.7 | No | ns | >0.9999 |
| WT:Sham vs. Chrdl1 KO:Stroke | -609.00 | -1227 to 8.703 | No | ns | 0.0540 |
| WT:Stroke vs. Chrdl1 KO:Sham | 1026.00 | 408.1 to 1643 | Yes | ** | 0.0018 |
| WT:Stroke vs. Chrdl1 KO:Stroke | 409.00 | -251.3 to 1069 | No | ns | 0.3562 |
| Chrdl1 KO:Sham vs. Chrdl1 KO:Stroke | -616.80 | -1234 to 0.8791 | No | ns | 0.0504 |

**Iba1 intensity x %area (contra)** Supplementary figure 4

**24 hours post-stroke**

| CONTRA | Mean | Std. Error of Mean | N |
|---|---|---|---|
| WT sham | 115.7 | 65.8 | 3 |
| WT stroke | 76.1 | 45.9 | 5 |
| Chrdl1 KO sham | 94.0 | 46.8 | 3 |
| Chrdl1 KO stroke | 108.2 | 79.4 | 5 |

| Šídák's multiple comparisons test | Predicted (LS) mean diff. | 95.00% CI of diff. | Below threshold? | Summary | Adjusted P Value |
|---|---|---|---|---|---|
| WT:Sham vs. WT:Stroke | 39.57 | -219.6 to 298.7 | No | ns | 0.9978 |
| WT:Sham vs. Chrdl1 KO:Sham | 21.66 | -268.1 to 311.4 | No | ns | >0.9999 |
| WT:Sham vs. Chrdl1 KO:Stroke | 7.49 | -251.7 to 266.6 | No | ns | >0.9999 |
| WT:Stroke vs. Chrdl1 KO:Sham | -17.91 | -277.0 to 241.2 | No | ns | >0.9999 |
| WT:Stroke vs. Chrdl1 KO:Stroke | -32.08 | -256.5 to 192.3 | No | ns | 0.9985 |
| Chrdl1 KO:Sham vs. Chrdl1 KO:Stroke | -14.18 | -273.3 to 245.0 | No | ns | >0.9999 |

**7 days post-stroke**

| CONTRA | Mean | Std. Error of Mean | N |
|---|---|---|---|
| WT sham | 27.1 | 7.5 | 4 |
| WT stroke | 30.5 | 15.0 | 3 |
| Chrdl1 KO sham | 14.2 | 5.0 | 4 |
| Chrdl1 KO stroke | 20.6 | 8.4 | 3 |

| Šídák's multiple comparisons test | Predicted (LS) mean diff. | 95.00% CI of diff. | Below threshold? | Summary | Adjusted P Value |
|---|---|---|---|---|---|
| WT:Sham vs. WT:Stroke | -3.40 | -44.66 to 37.86 | No | ns | >0.9999 |
| WT:Sham vs. Chrdl1 KO:Sham | 12.94 | -25.26 to 51.14 | No | ns | 0.8768 |
| WT:Sham vs. Chrdl1 KO:Stroke | 6.58 | -34.68 to 47.84 | No | ns | 0.9967 |
| WT:Stroke vs. Chrdl1 KO:Sham | 16.34 | -24.92 to 57.61 | No | ns | 0.7834 |
| WT:Stroke vs. Chrdl1 KO:Stroke | 9.98 | -34.13 to 54.09 | No | ns | 0.9796 |
| Chrdl1 KO:Sham vs. Chrdl1 KO:Stroke | -6.36 | -47.63 to 34.90 | No | ns | 0.9972 |

**Spine density (Ipsi)** Figure 5

**24 hours post-stroke**

| IPSI | protrusions/um Mean | Std. Error of Mean | N |
|---|---|---|---|
| WT sham | 1.17 | 0.08 | 3 |
| WT stroke | 0.50 | 0.11 | 5 |
| Chrdl1 KO sham | 1.10 | 0.15 | 3 |
| Chrdl1 KO stroke | 1.22 | 0.17 | 5 |

| Šídák's multiple comparisons test | Predicted (LS) mean diff. | 95.00% CI of diff. | Below threshold? | Summary | Adjusted P Value |
|---|---|---|---|---|---|
| Sham:WT vs. Sham:Chrdl1 KO | 0.07 | -0.6731 to 0.8124 | No | ns | 0.9999 |
| Sham:WT vs. Stroke:WT | 0.67 | 0.002999 to 1.332 | Yes | * | 0.0487 |
| Sham:WT vs. Stroke:Chrdl1 KO | -0.06 | -0.7228 to 0.6059 | No | ns | >0.9999 |
| Sham:Chrdl1 KO vs. Stroke:WT | 0.60 | -0.06667 to 1.262 | No | ns | 0.0883 |
| Sham:Chrdl1 KO vs. Stroke:Chrdl1 KO | -0.13 | -0.7925 to 0.5362 | No | ns | 0.9923 |
| Stroke:WT vs. Stroke:Chrdl1 KO | -0.73 | -1.301 to -0.1505 | Yes | * | 0.0113 |

**7 days post-stroke**

| IPSI | protrusions/um Mean | Std. Error of Mean | N |
|---|---|---|---|
| WT sham | 1.46 | 0.12 | 4 |
| WT stroke | 0.91 | 0.08 | 5 |
| Chrdl1 KO sham | 1.47 | 0.15 | 4 |
| Chrdl1 KO stroke | 1.25 | 0.08 | 5 |

| Šídák's multiple comparisons test | Predicted (LS) mean diff. | 95.00% CI of diff. | Below threshold? | Summary | Adjusted P Value |
|---|---|---|---|---|---|
| Sham:WT vs. Sham:Chrdl1 KO | -0.01 | -0.4845 to 0.4655 | No | ns | >0.9999 |
| Sham:WT vs. Stroke:WT | 0.55 | 0.09662 to 0.9979 | Yes | * | 0.0138 |
| Sham:WT vs. Stroke:Chrdl1 KO | 0.21 | -0.2418 to 0.6595 | No | ns | 0.6921 |
| Sham:Chrdl1 KO vs. Stroke:WT | 0.56 | 0.1061 to 1.007 | Yes | * | 0.0122 |
| Sham:Chrdl1 KO vs. Stroke:Chrdl1 KO | 0.22 | -0.2323 to 0.6690 | No | ns | 0.6501 |
| Stroke:WT vs. Stroke:Chrdl1 KO | -0.34 | -0.7633 to 0.08645 | No | ns | 0.1609 |

**Spine morphology (Ipsi)**

**24 hours post-stroke**

| IPSI | mature protrusions/um Mean | Std. Error of Mean | N |
|---|---|---|---|
| WT sham | 0.33 | 0.02 | 3 |
| WT stroke | 0.10 | 0.03 | 5 |
| Chrdl1 KO sham | 0.31 | 0.06 | 3 |
| Chrdl1 KO stroke | 0.33 | 0.04 | 5 |

| Tukey's multiple comparisons test | Mean Diff. | 95.00% CI of diff. | Below threshold? | Summary | Adjusted P Value |
|---|---|---|---|---|---|
| WT sham vs. WT stroke | 0.23 | 0.06139 to 0.3990 | Yes | ** | 0.0076 |
| WT sham vs. KO sham | 0.02 | -0.1687 to 0.2087 | No | ns | 0.9887 |
| WT sham vs. KO stroke | -0.01 | -0.1740 to 0.1636 | No | ns | 0.9997 |
| WT stroke vs. KO sham | -0.21 | -0.3790 to -0.04139 | Yes | * | 0.0140 |
| WT stroke vs. KO stroke | -0.24 | -0.3816 to -0.08920 | Yes | ** | 0.0022 |
| KO sham vs. KO stroke | -0.03 | -0.1940 to 0.1436 | No | ns | 0.9697 |

| IPSI | immature protrusions/um Mean | Std. Error of Mean | N |
|---|---|---|---|
| WT sham | 0.84 | 0.08 | 3 |
| WT stroke | 0.40 | 0.08 | 5 |
| Chrdl1 KO sham | 0.79 | 0.12 | 3 |
| Chrdl1 KO stroke | 0.89 | 0.13 | 5 |

| Tukey's multiple comparisons test | Mean Diff. | 95.00% CI of diff. | Below threshold? | Summary | Adjusted P Value |
|---|---|---|---|---|---|
| WT sham vs. WT stroke | 0.44 | -0.04634 to 0.9209 | No | ns | 0.0813 |
| WT sham vs. KO sham | 0.05 | -0.4907 to 0.5907 | No | ns | 0.9924 |
| WT sham vs. KO stroke | -0.05 | -0.5363 to 0.4309 | No | ns | 0.9877 |
| WT stroke vs. KO sham | -0.39 | -0.8709 to 0.09634 | No | ns | 0.1348 |
| WT stroke vs. KO stroke | -0.49 | -0.9088 to -0.07118 | Yes | * | 0.0207 |
| KO sham vs. KO stroke | -0.10 | -0.5863 to 0.3809 | No | ns | 0.9202 |

**7 days post-stroke**

| IPSI | mature protrusions/um Mean | Std. Error of Mean | N |
|---|---|---|---|
| WT sham | 0.28 | 0.06 | 4 |
| WT stroke | 0.20 | 0.01 | 5 |
| Chrdl1 KO sham | 0.31 | 0.08 | 4 |
| Chrdl1 KO stroke | 0.29 | 0.06 | 5 |

| Tukey's multiple comparisons test | Mean Diff. | 95.00% CI of diff. | Below threshold? | Summary | Adjusted P Value |
|---|---|---|---|---|---|
| WT sham vs. WT stroke | 0.08 | -0.1493 to 0.3052 | No | ns | 0.7535 |
| WT sham vs. KO sham | -0.03 | -0.2705 to 0.2085 | No | ns | 0.9811 |
| WT sham vs. KO stroke | -0.01 | -0.2409 to 0.2136 | No | ns | 0.9980 |
| WT stroke vs. KO sham | -0.11 | -0.3362 to 0.1183 | No | ns | 0.5232 |
| WT stroke vs. KO stroke | -0.09 | -0.3058 to 0.1226 | No | ns | 0.6115 |
| KO sham vs. KO stroke | 0.02 | -0.2099 to 0.2446 | No | ns | 0.9960 |

| IPSI | immature protrusions/um Mean | Std. Error of Mean | N |
|---|---|---|---|
| WT sham | 1.18 | 0.13 | 4 |
| WT stroke | 0.71 | 0.08 | 5 |
| Chrdl1 KO sham | 1.16 | 0.21 | 4 |
| Chrdl1 KO stroke | 0.96 | 0.02 | 5 |

| Tukey's multiple comparisons test | Mean Diff. | 95.00% CI of diff. | Below threshold? | Summary | Adjusted P Value |
|---|---|---|---|---|---|
| WT sham vs. WT stroke | 0.47 | -0.01906 to 0.9581 | No | ns | 0.0615 |
| WT sham vs. KO sham | 0.02 | -0.4937 to 0.5362 | No | ns | 0.9994 |
| WT sham vs. KO stroke | 0.22 | -0.2659 to 0.7113 | No | ns | 0.5631 |
| WT stroke vs. KO sham | -0.45 | -0.9368 to 0.04031 | No | ns | 0.0771 |
| WT stroke vs. KO stroke | -0.25 | -0.7074 to 0.2138 | No | ns | 0.4322 |
| KO sham vs. KO stroke | 0.20 | -0.2871 to 0.6900 | No | ns | 0.6377 |

**Spine density (contra)** Supplementary figure 5

**24 hours post-stroke**

protrusions/um

| CONTRA | Mean | Std. Error of Mean | N |
|---|---|---|---|
| WT sham | 1.26 | 0.26 | 3 |
| WT stroke | 1.18 | 0.14 | 5 |
| Chrdl1 KO sham | 1.44 | 0.36 | 3 |
| Chrdl1 KO stroke | 1.15 | 0.07 | 5 |

| Šídák's multiple comparisons test | Predicted (LS) mean diff. | 95.00% CI of diff. | Below threshold? | Summary | Adjusted P Value |
|---|---|---|---|---|---|
| Sham:WT vs. Sham:Chrdl1 KO | -0.18 | -1.139 to 0.7800 | No | ns | 0.9935 |
| Sham:WT vs. Stroke:WT | 0.09 | -0.7717 to 0.9444 | No | ns | 0.9998 |
| Sham:WT vs. Stroke:Chrdl1 KO | 0.11 | -0.7487 to 0.9674 | No | ns | 0.9992 |
| Sham:Chrdl1 KO vs. Stroke:WT | 0.27 | -0.5924 to 1.124 | No | ns | 0.9246 |
| Sham:Chrdl1 KO vs. Stroke:Chrdl1 KO | 0.29 | -0.5694 to 1.147 | No | ns | 0.8934 |
| Stroke:WT vs. Stroke:Chrdl1 KO | 0.02 | -0.7201 to 0.7661 | No | ns | >0.9999 |

**7 days post-stroke**

protrusions/um

| CONTRA | Mean | Std. Error of Mean | N |
|---|---|---|---|
| WT sham | 1.45 | 0.08 | 4 |
| WT stroke | 1.40 | 0.08 | 5 |
| Chrdl1 KO sham | 1.60 | 0.17 | 4 |
| Chrdl1 KO stroke | 1.29 | 0.11 | 5 |

| Šídák's multiple comparisons test | Predicted (LS) mean diff. | 95.00% CI of diff. | Below threshold? | Summary | Adjusted P Value |
|---|---|---|---|---|---|
| Sham:WT vs. Sham:Chrdl1 KO | -0.14 | -0.6569 to 0.3679 | No | ns | 0.9547 |
| Sham:WT vs. Stroke:WT | 0.05 | -0.4362 to 0.5359 | No | ns | 0.9998 |
| Sham:WT vs. Stroke:Chrdl1 KO | 0.16 | -0.3242 to 0.6479 | No | ns | 0.9061 |
| Sham:Chrdl1 KO vs. Stroke:WT | 0.19 | -0.2917 to 0.6804 | No | ns | 0.8098 |
| Sham:Chrdl1 KO vs. Stroke:Chrdl1 KO | 0.31 | -0.1797 to 0.7924 | No | ns | 0.3715 |
| Stroke:WT vs. Stroke:Chrdl1 KO | 0.11 | -0.3463 to 0.5703 | No | ns | 0.9771 |

**Spine morphology (contra)**

**24 hours post-stroke**

mature protrusions/um

| CONTRA | Mean | Std. Error of Mean | N |
|---|---|---|---|
| WT sham | 0.25 | 0.07 | 3 |
| WT stroke | 0.25 | 0.04 | 5 |
| Chrdl1 KO sham | 0.26 | 0.11 | 3 |
| Chrdl1 KO stroke | 0.27 | 0.01 | 5 |

| Tukey's multiple comparisons test | Mean Diff. | 95.00% CI of diff. | Below threshold? | Summary | Adjusted P Value |
|---|---|---|---|---|---|
| WT sham vs. WT stroke | 0.00 | -0.2370 to 0.2330 | No | ns | >0.9999 |
| WT sham vs. KO sham | -0.01 | -0.2754 to 0.2500 | No | ns | 0.9989 |
| WT sham vs. KO stroke | -0.03 | -0.2610 to 0.2090 | No | ns | 0.9872 |
| WT stroke vs. KO sham | -0.01 | -0.2456 to 0.2243 | No | ns | 0.9991 |
| WT stroke vs. KO stroke | -0.02 | -0.2275 to 0.1795 | No | ns | 0.9845 |
| KO sham vs. KO stroke | -0.01 | -0.2483 to 0.2216 | No | ns | 0.9982 |

immature protrusions/um

| CONTRA | Mean | Std. Error of Mean | N |
|---|---|---|---|
| WT sham | 1.02 | 0.21 | 3 |
| WT stroke | 0.93 | 0.12 | 5 |
| Chrdl1 KO sham | 1.18 | 0.25 | 3 |
| Chrdl1 KO stroke | 0.88 | 0.08 | 5 |

| Tukey's multiple comparisons test | Mean Diff. | 95.00% CI of diff. | Below threshold? | Summary | Adjusted P Value |
|---|---|---|---|---|---|
| WT sham vs. WT stroke | 0.09 | -0.5579 to 0.7331 | No | ns | 0.9769 |
| WT sham vs. KO sham | -0.17 | -0.8890 to 0.5543 | No | ns | 0.8995 |
| WT sham vs. KO stroke | 0.13 | -0.5107 to 0.7803 | No | ns | 0.9237 |
| WT stroke vs. KO sham | -0.25 | -0.9004 to 0.3905 | No | ns | 0.6543 |
| WT stroke vs. KO stroke | 0.05 | -0.5118 to 0.6062 | No | ns | 0.9942 |
| KO sham vs. KO stroke | 0.30 | -0.3433 to 0.9476 | No | ns | 0.5285 |

**7 days post-stroke**

mature protrusions/um

| CONTRA | Mean | Std. Error of Mean | N |
|---|---|---|---|
| WT sham | 0.33 | 0.05 | 4 |
| WT stroke | 0.28 | 0.03 | 5 |
| Chrdl1 KO sham | 0.32 | 0.01 | 4 |
| Chrdl1 KO stroke | 0.29 | 0.05 | 5 |

| Tukey's multiple comparisons test | Mean Diff. | 95.00% CI of diff. | Below threshold? | Summary | Adjusted P Value |
|---|---|---|---|---|---|
| WT sham vs. WT stroke | 0.05 | -0.1062 to 0.2132 | No | ns | 0.7662 |
| WT sham vs. KO sham | 0.01 | -0.1576 to 0.1791 | No | ns | 0.9976 |
| WT sham vs. KO stroke | 0.04 | -0.1172 to 0.2022 | No | ns | 0.8650 |
| WT stroke vs. KO sham | -0.04 | -0.2024 to 0.1169 | No | ns | 0.8630 |
| WT stroke vs. KO stroke | -0.01 | -0.1615 to 0.1395 | No | ns | 0.9965 |
| KO sham vs. KO stroke | 0.03 | -0.1279 to 0.1914 | No | ns | 0.9371 |

immature protrusions/um

| CONTRA | Mean | Std. Error of Mean | N |
|---|---|---|---|
| WT sham | 1.12 | 0.09 | 4 |
| WT stroke | 1.12 | 0.07 | 5 |
| Chrdl1 KO sham | 1.28 | 0.18 | 4 |
| Chrdl1 KO stroke | 0.95 | 0.13 | 5 |

| Tukey's multiple comparisons test | Mean Diff. | 95.00% CI of diff. | Below threshold? | Summary | Adjusted P Value |
|---|---|---|---|---|---|
| WT sham vs. WT stroke | 0.00 | -0.4938 to 0.4865 | No | ns | >0.9999 |
| WT sham vs. KO sham | -0.16 | -0.6721 to 0.3611 | No | ns | 0.8177 |
| WT sham vs. KO stroke | 0.17 | -0.3156 to 0.6647 | No | ns | 0.7323 |
| WT stroke vs. KO sham | -0.15 | -0.6420 to 0.3383 | No | ns | 0.8047 |
| WT stroke vs. KO stroke | 0.18 | -0.2839 to 0.6403 | No | ns | 0.6832 |
| KO sham vs. KO stroke | 0.33 | -0.1601 to 0.8202 | No | ns | 0.2497 |
